# Multi-strain phage induced clearance of bacterial infections

**DOI:** 10.1101/2024.09.07.611814

**Authors:** Jacopo Marchi, Chau Nguyen Ngoc Minh, Laurent Debarbieux, Joshua S. Weitz

**Author notes:** Electronic address (Jacopo Marchi). Electronic address - Former affiliation: School of Biological Sciences, Georgia Institute of Technology, Atlanta, GA, USA.

## Abstract

Bacteriophage (or ‘phage’ – viruses that infect and kill bacteria) are increasingly considered as a therapeutic alternative to treat antibiotic-resistant bacterial infections. However, bacteria can evolve resistance to phage, presenting a significant challenge to the near- and long-term success of phage therapeutics. Application of mixtures of multiple phage (i.e., ‘cocktails’) have been proposed to limit the emergence of phage-resistant bacterial mutants that could lead to therapeutic failure. Here, we combine theory and computational models of *in vivo* phage therapy to study the efficacy of a phage cocktail, composed of two complementary phages motivated by the example of *Pseudomonas aeruginosa* facing two phages that exploit different surface receptors, LUZ19v and PAK P1. As confirmed in a Luria-Delbrück fluctuation test, this motivating example serves as a model for instances where bacteria are extremely unlikely to develop simultaneous resistance mutations against both phages. We then quantify therapeutic outcomes given single- or double-phage treatment models, as a function of phage traits and host immune strength. Building upon prior work showing monophage therapy efficacy in immunocompetent hosts, here we show that phage cocktails comprised of phage targeting independent bacterial receptors can improve treatment outcome in immunocompromised hosts and reduce the chance that pathogens simultaneously evolve resistance against phage combinations. The finding of phage cocktail efficacy is qualitatively robust to differences in virus-bacteria interactions and host immune dynamics. Altogether, the combined use of theory and computational analysis highlights the influence of viral life history traits and receptor complementarity when designing and deploying phage cocktails in immunocompetent and immunocompromised hosts.

## I. INTRODUCTION

Bacteriophage (phage, *i.e*. viruses that infect bacteria) are the most abundant organisms on the planet[1, 2]. Bacteria have evolved a myriad of defense mechanisms to tolerate, counter, and resist infections [3, 4]. Likewise, phages have co-evolved with bacteria for billions of years, catalyzing the emergence of a myriad of phage variants [5–8]. Diverse phage represent a largely untapped, therapeutic reservoir for treatment of antibiotic-resistant bacteria [9–12]. The therapeutic application of lytic phage is gaining interest as a viable treatment in alternative to antibiotics [13], capitalizing on the evolutionary dynamics of phages to combat bacterial pathogens that have outpaced traditional treatment options. Unlike antibiotics, which exert selective pressure driving the emergence of resistance, phage possess the inherent ability to evolve alongside bacteria, and can potentially adapt to overcome resistance mechanisms and maintain their efficacy over time [9]. Although evolution of phage resistance in bacteria is feasible, the widespread application of phage therapy is unlikely to catalyze broad spectrum resistance, given the high host specificity of phage [14].

In practice, phage therapy has shown significant potential in *in vivo* studies [15, 16] and in human treatment in compassionate cases [17, 18] and clinical trials [19]. The study [18] demonstrated that a personalized phage cocktail successfully treated a life-threatening, multidrug-resistant *Acinetobacter baumannii* infection in a critically ill patient, where conventional antibiotics had failed. However, there have been cases where therapeutic phage were not shown to be significantly more effective in controlling bacterial infections than standard procedures of care [20, 21]. In the authors found that a phage cocktail was safe and well-tolerated in treating *Pseudomonas aeruginosa* burn wound infections, but it did not demonstrate a significant improvement in bacterial load reduction or wound healing compared to standard treatment. The mixed results of clinical trials call for a thorough identification of the factors leading to success or failure of phage therapy [22].

To address this gap, several studies have integrated *in vitro* and *in vivo* experiments, as well as *in silico* computational models, into the design and assessment of phage therapy [23]. These studies have advanced the quantitative understanding of parameters impacting phage therapy outcomes, such as the pathogen strain, infected tissue, and chosen delivery method for the therapeutic phage [24]. A key factor modulating the outcome of *in vivo* phage therapy is the mammalian host immune response. For example, combined use of theory and *in vivo* application of phage in immunomodulated murine hosts revealed that phage and neutrophils work synergistically to eliminate *P. aeruginosa* and prevent the onset of fatal acute pneumonia [16]. In immunophage synergy, phage rapidly infect and lyse susceptible *P. aeruginosa* cells while neutrophils clear both susceptible and subpopulations of phage-resistance cells. However, not all phage and immune interactions may lead to positive impacts on therapy. For instance, alveolar macrophages can reduce the density of circulating phage, potentially jeopardizing phage therapeutic treatment efficacy [25]. When immune systems are compromised [26] or limit phage-induced clearance of target bacteria [25], the proliferation of phage-resistance mutants can lead to therapeutic failure [16].

Indeed phage resistance represents one of the major challenges to phage therapy success. The emergence of bacteria resistant to virulent phage reduces the ability of phage to contain an infection [27–29]. There are multiple approaches to overcome the evolution and proliferation of phage resistant bacteria. First, phage can be trained via asymmetric evolutionary training to target evolved bacteria that are resistance to the original, therapeutic phage [30]. Subsequent application of trained phage can limit the emergence of phage-resistant bacteria, improving therapeutic efficacy. Alternatively, phage may be used in combination with small molecules (e.g., antibiotics) that limit the potential for bacterial evolution. In one well-studied case, phage OMKO1 targets antibiotic efflux pumps within Pa; hence joint use of phage OMKO1 and antibiotics can lead to therapeutic success [31]. Finally, there may be cases where phage target receptor sites such that evolution of phage resistance comes with significant fitness costs in an in vivo context [9, 32–37]. All of these examples serve to illustrate the general rule that decreasing the scope of phage-escape bacterial mutants and/or increasing the costs of phage-resistant mutations increase the efficacy of phage therapy. In the same spirit, the combined use of multiple phages in a cocktail that infect distinct receptors, makes it harder for bacteria to acquire resistance against all phages at the same time [38–41]. Cocktails with phages targeting different cell receptors have higher efficacy against *P. aeruginosa* [42]. These evolutionary principles can be combined in cocktails that comprise trained phage, which can efficiently control *P. aeruginosa* populations both *in vitro* and in mice lung infections [43].

In this work we focus on evaluating the therapeutic efficacy of phage cocktails, which leverage complementary adsorption paths to infect target bacteria. We combine computational and analytical treatment of single- and double-phage therapy models, informed by Luria-Delbrück (LD) fluctuation tests exposing *P. aeruginosa* to phages LUZ19v and PAK P1, to address whether phage cocktails can help restore therapy success in immunocompromised hosts in face of phage resistance. We then extend our analysis to include *in vivo* immune system dynamics and structured phage-bacteria interactions, mapping the quantitative conditions for treatment success as a function of the immune state and therapeutic phages effectiveness. Comparing our exploration with the parameters extracted in [16] we note that the quantitative adsorption rate of any of the therapeutic phages may have drastic effects on the treatment outcomes jeopardizing the benefits of phage cocktails, highlighting the importance of selecting a combination of efficient phages against the pathogenic strains especially in hosts with a compromised immune system.

## II. METHODS

### A. Fluctuation test to infer the probability of resistance mutations of *P. aeruginosa* against two phages

We perform a LD fluctuation test to assess the likelihood that *P. aeruginosa* colonies randomly develop a resistance mutation against either phage PAK P1 or LUZ19v or against both phages at the same time. For each phage, we grow overnight 360 independent populations of strain PAK-lumi. 300 of these are plated against either PAK P1 or LUZ19v to count the number of resistant mutants, whereas the remaining 60 are used as control to count the CFUs in the absence of phage. To measure resistance against both phages simultaneously we grow 150 independent colonies and plate them against a mixture of PAK P1 and LUZ19v. More details on the fluctuation test protocol are in SI Sec. II. Then we use the resistant mutants counts to infer the probability at which cells develop resistance to PAK P1, LUZ19v or both, within one duplication. We run the inference through the web-tool bz-rates [44], which learns the model parameters via the Generating Function estimator from [45] (see SI Sec. III for more details on the inference).

### B. Mathematical models of phage therapy

#### 1. Summary

We study two different models of single- and double-phage therapy of bacterial infections in immunomodulated hosts leading to four different scenarios. Figure 1 sketches the models ingredients both in terms of ecological dynamics (panel A) as well as evolutionary multi-strain phage-bacteria interactions (panel B). We start from a simple model of a lung infection, mimicking a signaling deficient immune system as in the case of myeloid differentiation primary response gene 88-deficient mice (MyD88^*−/−*^), where the immune system is static as neutrophils are not recruited in the lungs [16] (Figure 1, A_1_). We first analyze a single-phage treatment model (Figure 1, B_1_), then we add a second phage to the cocktail to address its impact on treatment outcome (Figure 1, B_2_ top). Third, switching back to a single-phage therapy, we include more complex ecological dynamics to account for a responsive immune system, still modulated by its carrying capacity (limiting the availability of immune cells within the infected tissue). In this second single-phage treatment model we include intermediate stages of infected bacteria and add a non-linearity in phage infections to mimic a Michaelis-Menten phage-bacteria binding reaction (Figure 1, A_2_). Finally we study the effect of a phage cocktail with the inclusion of these new model ingredients, and explore the role of evolutionary relations between the combined therapeutic phages (Figure 1, B_2_ bottom). We map the phage therapy outcomes as a function of host immune strength and phage efficacy, depending on the different components of these four models.

**FIG. 1:**
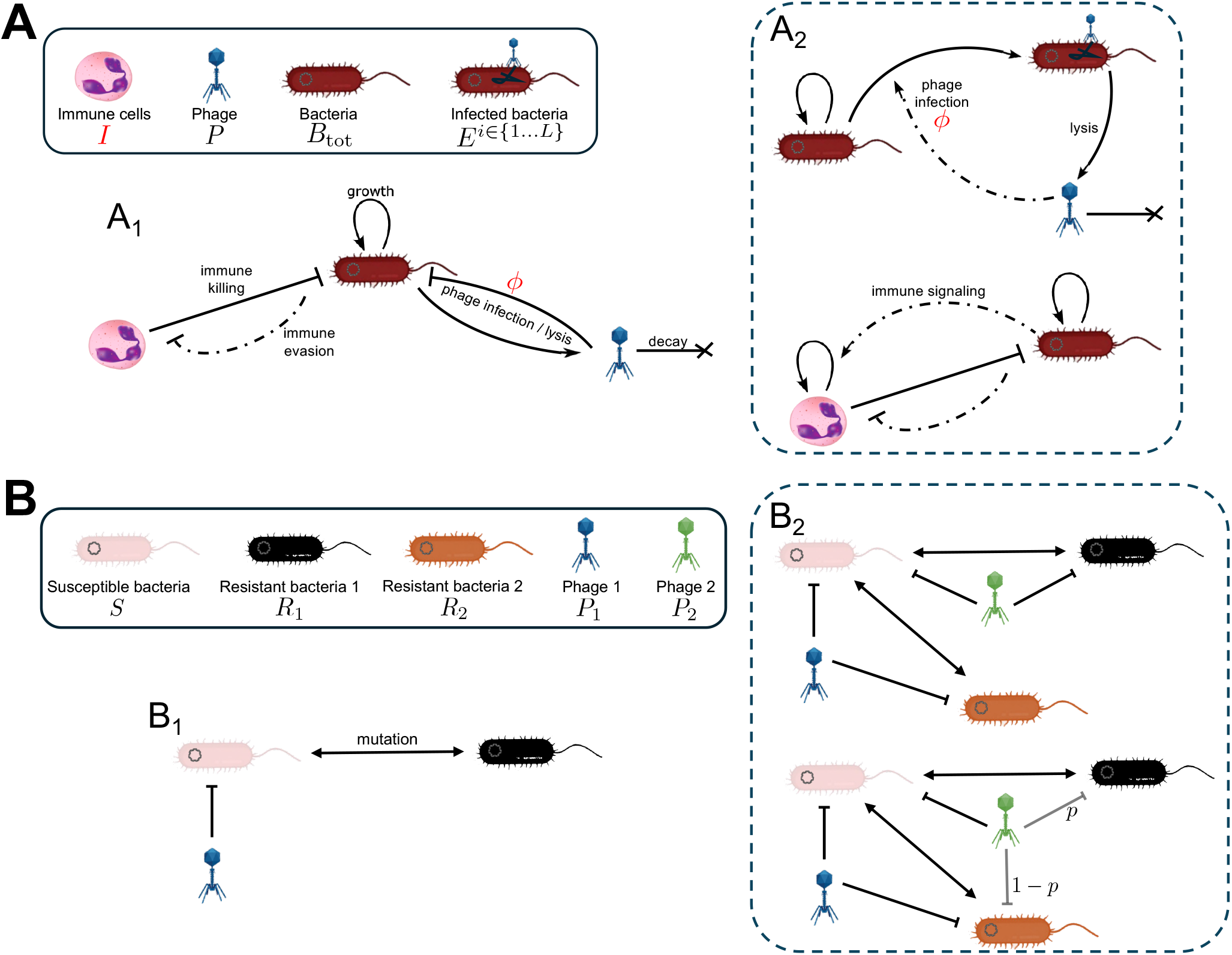
Schematic of mathematical model of phage therapy. A) Ecological interactions between bacteria, phage and immune system (graphical and mathematical symbols in the legend box). Subpanel A_1_ shows the interactions in the simple phage therapy model in a signaling deficient host introduced in Section II B 2, whereas Subpanel A_2_ sketches the addition of more complex dynamics in the model introduced in Section II B 4, namely a structured bacteria population including stages of phage infection and the active recruitment of the host immune cells. Panel B) gathers the cross-infection and mutation networks between specific phage and bacteria strains (listed in the legend box) in the different treatment models. Subpanel B_1_ sketches the single-phage treatment structure used in Sections II B 2 and II B 4, whereas B_2_ summarizes the cross-infection structure in the phage-cocktail treatment models introduced in Sections II B 3 and II B 5 (top and bottom respectively). The main model parameters studied in this work, the immune strength *I* and the phage adsorption rate *ϕ*, are colored in red. Arrow types encode the different kind of dynamical interactions (growth, killing, decay, mutations), with dashed-dotted lines indicating saturating nonlinear rates of the form 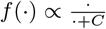.

#### 2. Infection model of interacting phage, multi-strain bacteria and non-responsive host immune system

The following ordinary differential equation model represents how phage *P* interacts with susceptible bacteria *S* and phage-resistant bacteria *R*, under the action of immune cells *I* during infection of an immunodeficient host:

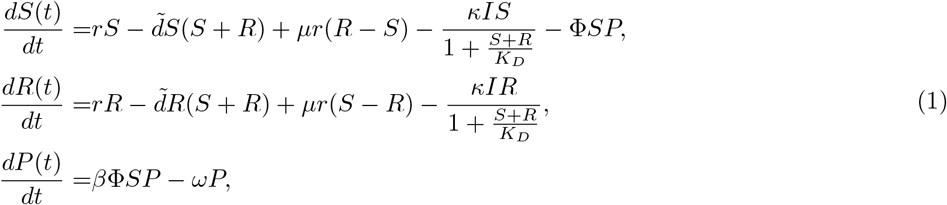

This model includes the main ingredients pinpointed in early theoretical studies of phage-bacteria interactions *in vitro* [47] and in the context of phage therapy [48]. Bacteria undergo logistic growth at maximum rate *r* and with density driven death rate 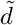, equivalent to a carrying capacity term 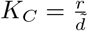. They mutate between the susceptible and resistant types, and the susceptible type can be infected by phage with adsorption rate *ϕ*. At the same time phage lyse susceptible bacteria producing *β* virions per lysis event, and decay at a constant rate *ω*. We build on previous models focusing on acute infections [16, 26], therefore we only include the action of innate immune effector cells *I*, mainly representing the action of neutrophils. In this first model we take *I* as a constant mimicking an immunodeficient host unable to recruit more immune cells other than a fixed baseline as in the case of myeloid differentiation primary response gene 88-deficient mice (MyD88^*−/−*^), as proposed in [16]. Immune cells target both bacteria types through a saturating function, as bacteria evade the immune action at high population densities compared to an evasion threshold *K*_*D*_, consistent with experiments showing that pathogens at high density can activate defenses against the immune system [49, 50] through physical shielding [51] or expression of virulence factors [52]. This modeling ingredient, included in [53] to explain the saturation in granulocytes action on *P. aeruginosa* during mice thigh infections and then adapted to the context of phage therapy [26], was shown to be essential to explain the synergy between phage and immune system in clearing bacteria infections [16, 26]. Therefore, model (1) represents a baseline model of immunophage synergy extending [26] to enable assessment of phage-resistant bacteria on in vivo dynamics given modulation of immune strength (*I*) and phage life history traits (here, primarily explored through variation in *ϕ*). Table I reports the parameters meaning and values.

**TABLE I:**
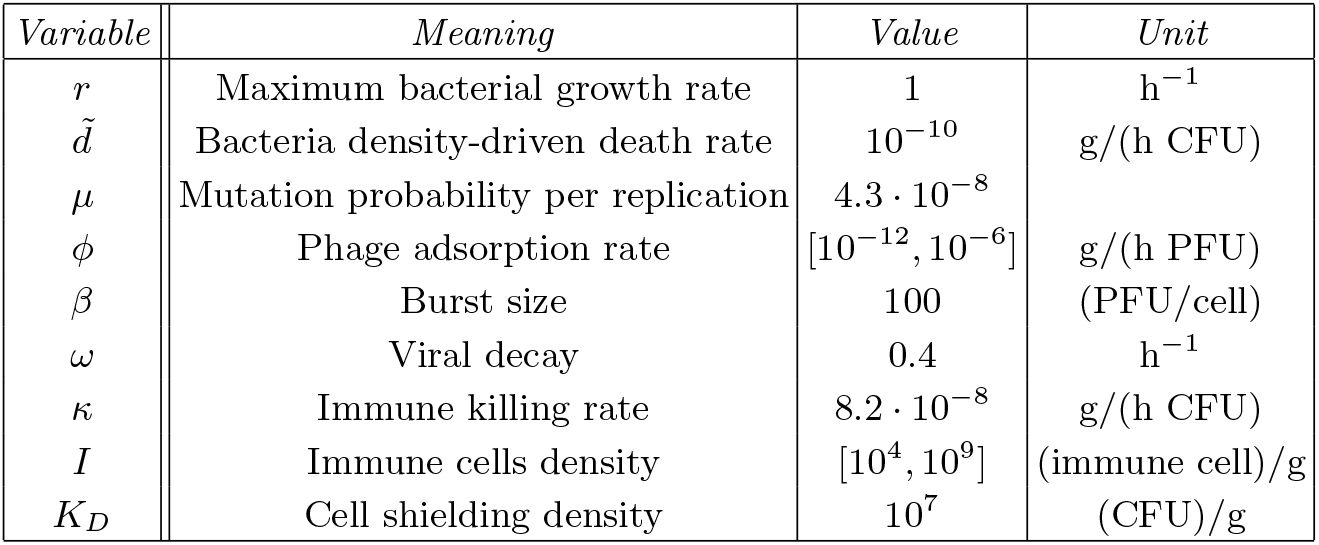
Model parameters. Most values are taken from [26]. The mutation rate is inferred from the fluctuation test (Section II A). For the viral decay *ω* we elect a mid-value between the parameter reported in [26, 46] (1 h^*−*1^) and the one inferred in [16] (0.07 h^*−*1^).

| *Variable* | *Meaning* | *Value* | *Unit* |
| --- | --- | --- | --- |
| $r$ | Maximum bacterial growth rate | 1 | $\text{h}^{-1}$ |
| $\tilde{d}$ | Bacteria density-driven death rate | $10^{-10}$ | $\text{g}/(\text{h CFU})$ |
| $\mu$ | Mutation probability per replication | $4.3 \cdot 10^{-8}$ | |
| $\phi$ | Phage adsorption rate | $[10^{-12}, 10^{-6}]$ | $\text{g}/(\text{h PFU})$ |
| $\beta$ | Burst size | 100 | $(\text{PFU}/\text{cell})$ |
| $\omega$ | Viral decay | 0.4 | $\text{h}^{-1}$ |
| $\kappa$ | Immune killing rate | $8.2 \cdot 10^{-8}$ | $\text{g}/(\text{h CFU})$ |
| $I$ | Immune cells density | $[10^4, 10^9]$ | $(\text{immune cell})/\text{g}$ |
| $K_D$ | Cell shielding density | $10^7$ | $(\text{CFU})/\text{g}$ |

#### 3. Model of a phage combination therapy against multiple pathogenic bacteria strains

In the second model we add a second phage to the infection treatment, now composed of a combination of two phages *P*_1_ and *P*_2_ (Figure 1, B_2_ top). The pathogen types include the wild-type infecting bacteria *S*, susceptible to both phages, which can in turn mutate to a type *R*_1_ resistant to *P*_1_ or to a type *R*_2_ resistant to *P*_2_ :

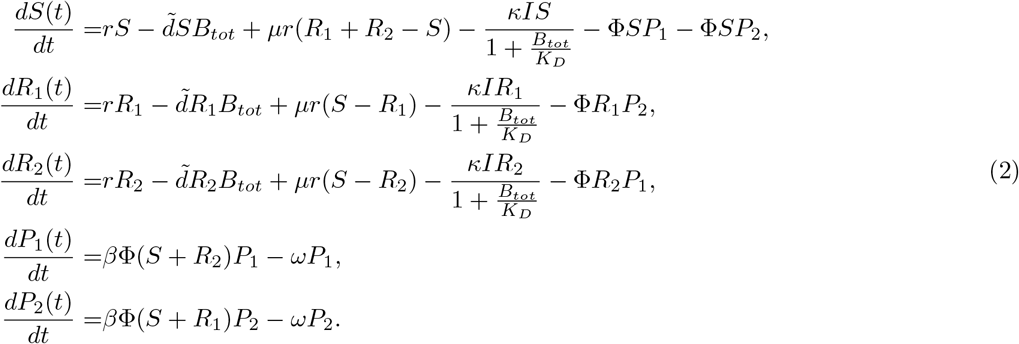

We initially assume that the two phages have the same adsorption rate *ϕ*, which we vary together with the immune strength *I* to evaluate therapeutic outcomes when using a phage cocktail. Importantly, Model (2) assumes that bacteria cannot develop resistance to both phages at the same time within an infection timescale, which was suggested in several studies proposing cocktail of phages that infect bacteria through different routes [41, 42, 54]. We also assume that there is no cross-infectivity between the two phages with respect to the resistant types. The susceptible bacteria mutate equally to either resistant type (and viceversa), but resistant types do not mutate directly among themselves – hence we ignore the near-term chance of the emergence of a double-phage resistant mutant.

#### 4. Structured model for single-phage treatment of in vivo infections with a modulated responsive immune system

We extend the single phage therapy model Eq. (1) to include a realistic *in vivo* innate immune response and more complex phage infection dynamics [16] (Figure 1, A_2_):

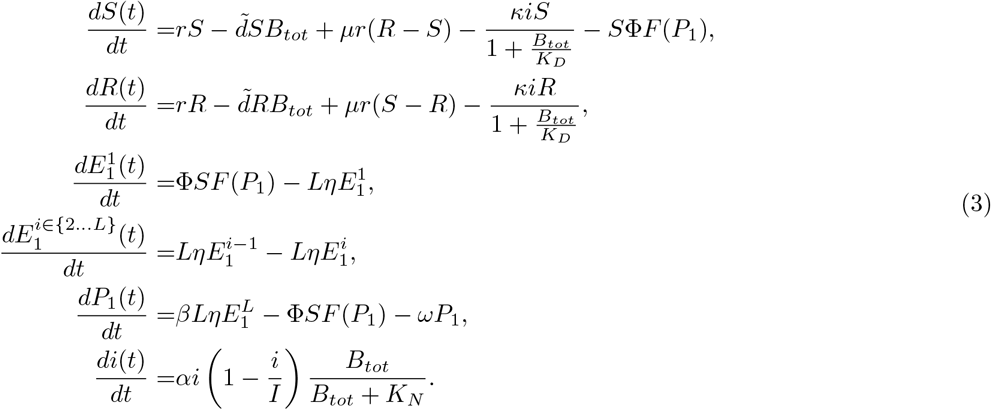

The innate immune cells *i* are recruited in the lungs at a maximum rate per capita *α* until they reach a saturation density *I* (immune capacity), which is one of the parameters we vary in this study to modulate hosts immune responses. Immune cells are recruited proportionally to a saturating function of the total density of bacteria *B*_*tot*_ with half saturation constant *K*_*N*_. These saturation profiles in immune recruitment within infected tissues are supported by empirical evidence that neutrophils can only be produced up to a limit [59], and that innate immune responses saturate as a function of bacterial loads [53], as already mentioned in the previous section.

Additionally, we structure the bacteria population introducing *L* infected bacteria stages, 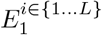, between phage adsorption and lysis to model a finite phage infection time distributed as an Erlang distribution with average 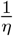, an approach applied in several models of epidemiological dynamics and phage-bacteria interactions [57, 60, 61]. Finally, as proposed in [16], we add a saturation term in phage adsorption 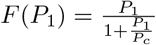, as if phage adsorbed into bacteria through Michaelis-Menten reaction kinetics [62, 63], producing a phage-adsorption profile similar to Monod growth [64]. Table II provides a list of parameter definitions and their values.

**TABLE II:** Model parameters. Most values are taken from [16]. The mutation rate is inferred via a fluctuation test (Section II). The lysis rate 2 h^*−*1^ is compatible with reported lysis times of phages infecting *P. aeruginosa* [55, 56]. The number of infected stages *L* is the same as several previous models of phage-bacteria dynamics [57, 58].

| *Variable* | *Meaning* | *Value* | *Unit* |
| --- | --- | --- | --- |
| $r$ | Maximum bacterial growth rate | 0.75 | $\text{h}^{-1}$ |
| $\tilde{d}$ | Bacteria density-driven death rate | $7.5 \cdot 10^{-9}$ | $\text{g}/(\text{h CFU})$ |
| $\mu$ | Mutation probability per replication | $4.3 \cdot 10^{-8}$ | |
| $\phi$ | Phage adsorption rate | $[10^{-9}, 10^{-6}]$ | $\text{g}/(\text{h PFU})$ |
| $P_c$ | Phage adsorption saturation density | $1.5 \cdot 10^7$ | $(\text{PFU})/\text{g}$ |
| $\eta$ | Lysis rate | 2 | $\text{h}^{-1}$ |
| $L$ | Number of infected stages | 10 | |
| $\beta$ | Burst size | 100 | $(\text{PFU}/\text{cell})$ |
| $\omega$ | Viral decay | 0.07 | $\text{h}^{-1}$ |
| $\kappa$ | Immune killing rate | $8.2 \cdot 10^{-8}$ | $\text{g}/(\text{h CFU})$ |
| $i$ | Immune cells capacity | $[10^4, 10^8]$ | $(\text{immune cell})/\text{g}$ |
| $K_D$ | Bacteria shielding density from immune action | $4.1 \cdot 10^7$ | $(\text{CFU})/\text{g}$ |
| $K_N$ | Bacteria density when immune recruitment is half its maximum | $10^7$ | $(\text{CFU})/\text{g}$ |
| $\alpha$ | Maximum immune cells recruitment rate | 0.97 | $\text{h}^{-1}$ |

#### 5. Multi-phage treatment model, modulating the evolutionary interactions between bacteria and phage strains during infections

Finally, we develop a multi-phage therapy model composed of two phages *P*_1_ and *P*_2_, in the presence of the new modeling ingredients presented in the previous section. As in eq. (2), bacteria susceptible to both phages, *S*, can mutate to a type *R*_1_ resistant to *P*_1_ or to a type *R*_2_ that can be infected by *P*_1_. Compared to section II B 3, now we introduce a parameter *p* continuously modulating the evolutionary relations between the two phages and the target bacteria, in an asymmetric fashion. *P*_2_ can either infect *R*_1_ with probability *p*, or *R*_2_ with probability 1*− p*. So when *p* = 1 the two phages are complementary in infecting each other’s resistant type with rate *ϕ*, and *R*_*i*_ truly has the meaning of bacteria resistant to phage *i*. When *p* = 0 the two phages are phenotypically equivalent in the sense that they target the same set of bacteria, as if resistance would arise to both at the same time. Intermediate values of *p* mimic a generalist phage *P*_2_ capable of infecting either bacterium, but with lower specificity to either type since the sum of the effective adsorption rates to all bacteria is always *ϕ*. The equations for this model are:

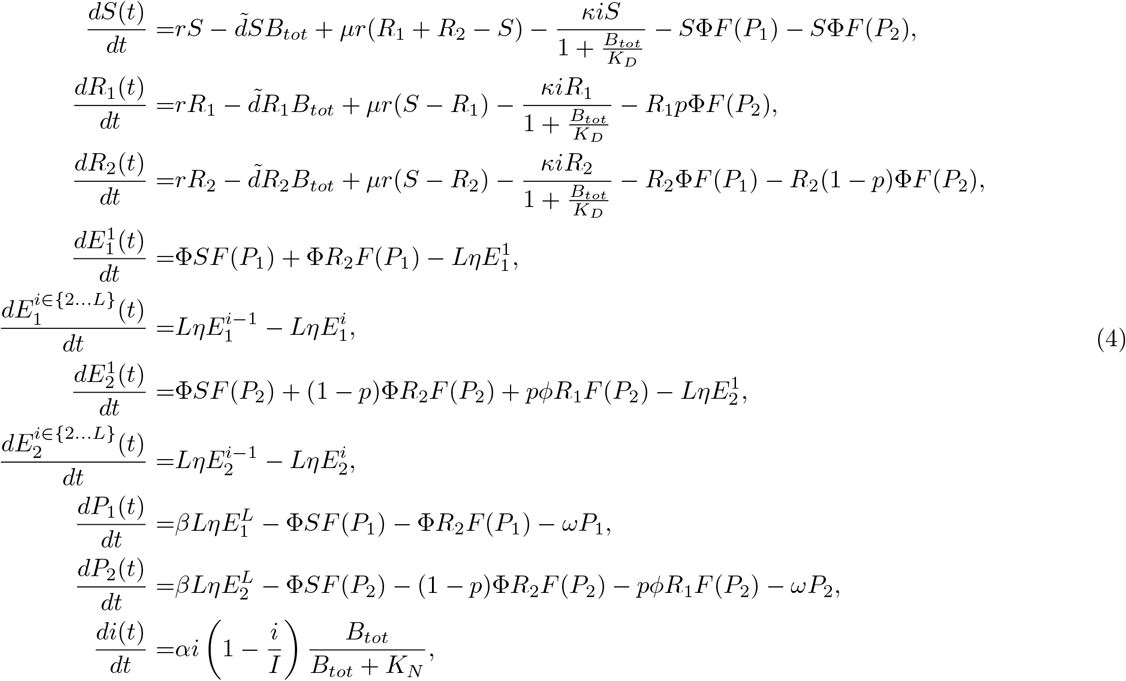

where 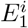 denotes a bacterium infected by *P*_1_ (at infection stage *i*), and 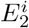 denotes a bacterium infected by *P*_2_. We start by considering phages with the same life history traits, and then relax this assumption studying a case where *P*_1_ adsorbs at a rate *ϕ*_1_ while *P*_2_ adsorbs at a rate *ϕ*_2_. In this case we study a scenario where 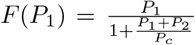,mimicking apparent competition between the two phages, as if they bind to the same “substrate” or surface receptor, or if they shared common molecular machinery in order to inject genetic material into the cell.

## III. RESULTS

### A. Condition for single-phage therapy success in the face of phage resistance and non-responsive immunity

We start by analyzing the single-phage model presented in Section II B 2. In the absence of phage, the model admits a stable fix point 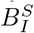 and an unstable one 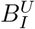 for bacteria densities. After the addition of phage, whenever bacteria are driven below the infectious dose 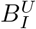 then the immune system is able to clear the infection, producing a successful treatment with phages and immune system working in synergy to control the pathogens [16, 26]. Neglecting the emergence of phage-resistant bacteria, the phage induced bacteria steady state becomes unstable when 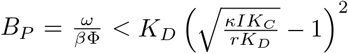 (derived in [26]). In terms of immune strength, this means that for immunophage synergy to function then the host innate immunity needs to be bigger than a threshold *I*_0_:

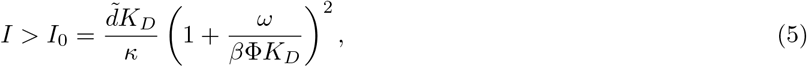

which in the absence of phage resistance would be a sufficient condition for therapeutic success.

In contrast, if phage-resistant bacteria are present in the population, then the phage-induced dynamical instability is not enough to drive bacteria down as the phage resistant bacteria grow to an immune-dependent stable density (see SI Sec. III B). The mathematical condition 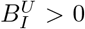 becomes necessary for therapy success together with Eq. (5), so that the bacteria extinction steady state *B*^*∗*^ = 0 becomes stable. In this case, phage can drive the susceptible bacteria below 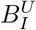 before the resistant bacteria proliferate from rare to densities above this threshold. When bacteria densities are depleted below 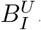, the immune system can clear the remainder of susceptible and resistant bacteria in the host. This mechanism restores the ability of phage and the host immune system to synergistically clear an infection in the presence of phage resistant bacteria, reinforcing phage-immune synergy against multiple strains of bacteria. Fig. 2 shows how, with 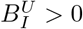, crossing the threshold *I > I*_0_ drives a transition from therapeutic failure to success as phage drive the susceptible bacteria below 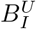 before the resistant type grows enough, at which point the immune system can clear both phage-susceptible and phage-resistant bacteria.

**FIG. 2:**
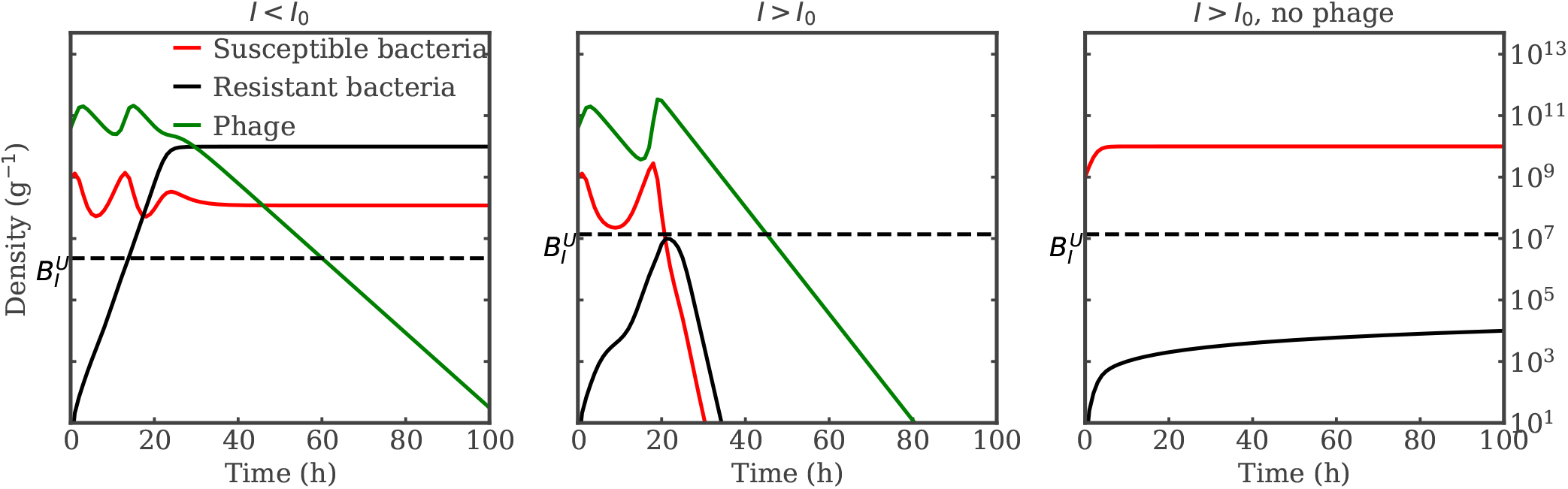
Phage-immune synergy in face of phage resistance. Simulated dynamics for phage, susceptible and resistant bacteria, when the immune system strength is below *I*_0_ (left panel) or above (center, right panels), with phage (left, center) or without (right). Increasing the immune system strength above *I*_0_, with 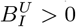, phage can drive susceptible bacteria below 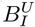 before the resistant type grows, driving a transition from therapy failure to success (left to center panel). This is a signature of phage-immune synergy as the infection would persist without phage (right). Simulation parameters are reported in Table I with *ϕ* = 10^11^ g/(h PFU), *I* = 1.5 *·* 10^7^ and 2.9 *·* 10^7^ cells/g respectively in the left and the other two panels.

Next, we assume that the immune evasion threshold *K*_*D*_ is lower than bacteria carrying capacity *K*_*C*_ and that the immune system is well below the strength where it would clear the infection on its own (and hence no therapy would be necessary). In this case we can derive the immune strength attaining 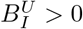, necessary for immunophage synergy (see SI Sec. III B for the derivation):

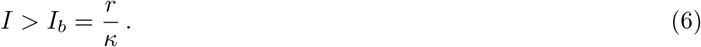

If phage drive the susceptible bacteria below 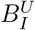 fast enough, the simultaneous satisfaction of the conditions in Eqs. (5) and (6), or more compactly

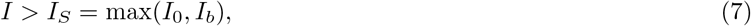

yields phage therapy success. This condition summarizes the necessary host immune strength as a function of phage life history traits, so that the application of a single phage clears the infection in an immunomodulated host, overcoming the evolutionary challenge posed by bacteria developing resistance to phage. The intuitive explanation of this generalization of immunophage synergy is as follows. First, the immune system needs to be strong enough so phage therapy can control susceptible bacteria, represented by the condition in Eq. (5). Second, resistant bacteria must be controllable at relatively low densities by the immune system, Eq. (6). Combining the two we recover the complete success condition in Eq. (7).

We test our analytic results through several simulations of the model in Eq. (1) varying the immune system strength *I* and the phage adsorption rate *ϕ*. Fig 3 shows the phase diagram for the density of bacteria after 200 hrs postinfection, comparable to in vivo experiments, averaged over the last day (details in SI Sec. III A). In this case bacteria are either eliminated in a successful treatment, or phage-resistant types grow to an immune system-dependent stable density and therapy fails. As predicted by Eq. (7), and confirmed by the numerical results, the immune system needs to be strong enough for a single-phage therapy to be successful and phage with a high adsorption rate can lead to therapy success in weaker hosts (lower *I*).

**FIG. 3:**
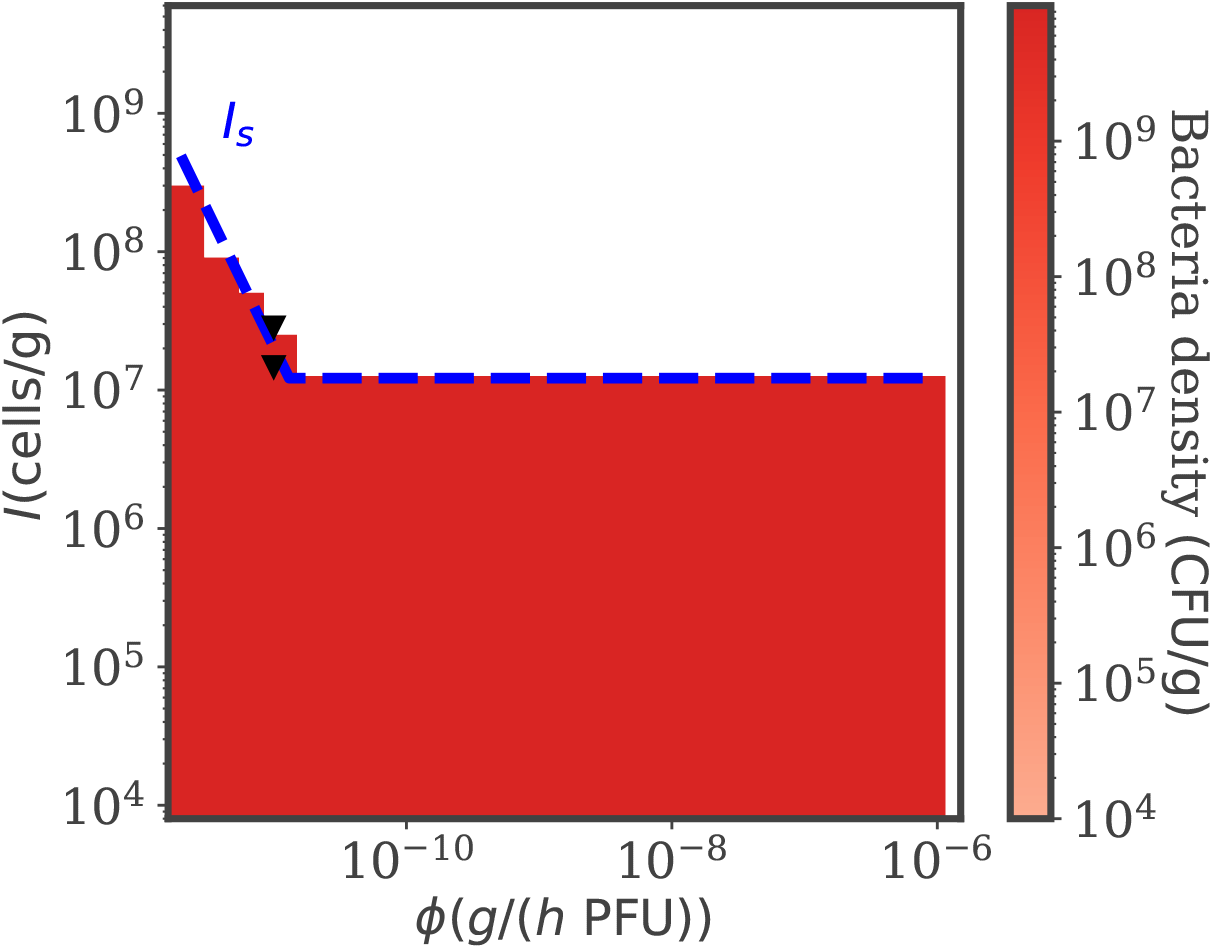
Bacteria density as a function of immune strength and phage adsorption rate. Numerical simulations of the model in Eq. (1) varying *I* and *ϕ*. The color map represents the density of bacteria in the last part of the numerical simulations. Single-phage therapy works such that bacteria are driven to elimination with a strong enough immune system (denoted in the white region). Efficient phage (higher *ϕ*) broaden the therapy success conditions. The analytic condition in Eq. (7) (blue dashed line) accurately predicts the therapy outcome transition. The two black triangles correspond to the left and central panel in Fig. 2. Simulation parameters are reported in Table I.

### B. Phage cocktails can improve therapy efficacy and overcome phage resistance in immunocompromised hosts

Immunophage synergy can eliminate a population of phage-resistant and phage-susceptible bacteria insofar as the immune system is sufficiently strong given the life history traits of the therapeutic phage (as summarized in Eqs.(5) and (6)). Likewise, monophage therapy can fail due to the proliferation of phage-resistant bacteria. Hence, we next explore how phage cocktails can potentially expand the range of immunocompromised hosts in which phage therapy is effective. To do so, we simulate Model (2), which includes two phages that can target the susceptible bacteria as well as one additional bacterium (i.e., each of the resistant bacteria is only resistant to one of the two phages in the cocktails). This complementarity with respect to resistance reflects that, in our *in vitro* experiment growing 150 separate populations of *P. aeruginosa*, we found no mutant conferring simultaneous resistance against a cocktail of phages PAK P1 and LUZ19v. Hence we assume that the emergence of simultaneous resistance against both phages from a background of susceptible bacteria is negligible.

Figure 4 A shows the expanded region of immunophage synergy given a phage cocktail via numerical simulations of model (2). The addition of a second phage can restore therapeutic success in immunocompromised hosts, provided phages have sufficiently effective life history traits (here, we focus on variation in the adsorption rate *ϕ*). This is expected as the added phage will lyse the bacteria resistant to the other one, as shown in the population dynamics in Figure 4C,D, which show how bacteria strains evolve under the selection imposed by phages eventually leading to extinction. Even so, if phages have low adsorption rate then they will not clear the infection (*e.g*. see dynamics in Figure 4B).

**FIG. 4:**
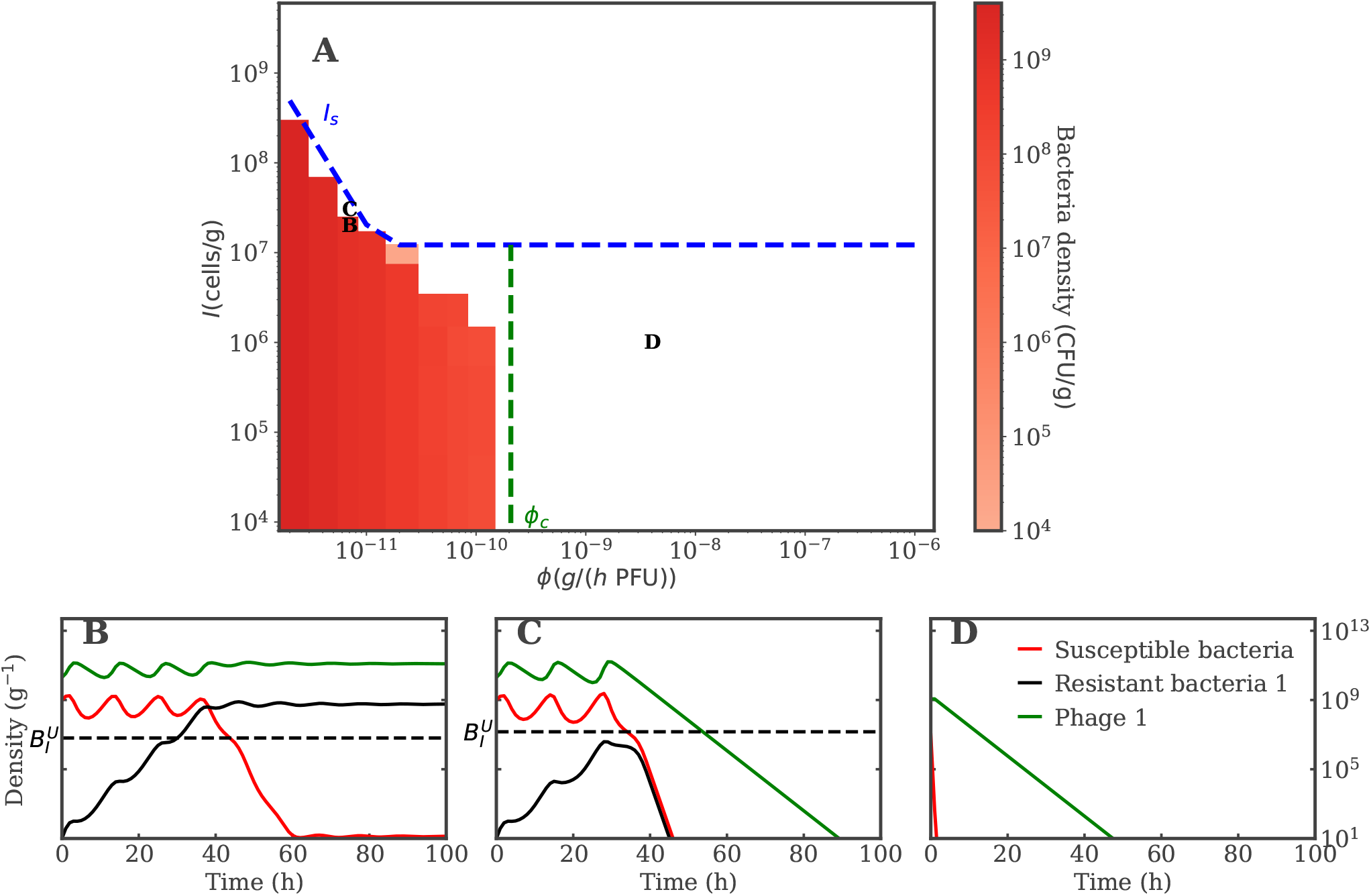
Phage cocktail therapy succeeds for a wide parameters range. A) Numerical simulations of the model in Eq. (2) varying *I* and *ϕ*. The color map represents the density of bacteria in the last part of the numerical simulations. Adding a second phage clears the infection for a much wider parameter range compared to single-phage treatment, if phages have high enough *ϕ*, as highlighted by the comparison with the transition for single-phage therapy success in Eq. (7) (blue dashed line). The vertical dashed green line represents the value of *ϕ*_*c*_ obtained solving Eq. (8) with Ω = 10 The black triangles correspond to the parameters yielding the population dynamics in panels B),C),D) that show the evolution of phage and bacteria strains during the simulated infection. Simulation parameters are reported in Table I.

We propose a heuristic derivation of the phase space of immunophage synergy, as detailed in SI Sec. III C. The result predicts that without the immune system (*I* = 0) phages drive bacteria to the extinction threshold *T* (1 bacterial cell in the lungs) for *ϕ > ϕ*_*c*_, where *ϕ*_*c*_ satisfies the transcendental equation:

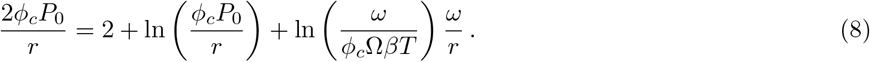

Here *P*_0_ is the initial phage density. Ω is a numerical factor ensuring bacteria exponential decay before extinction (see SI Sec. III C), which needs to satisfy Ω *≫* 1 and 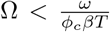. Ω affects the equation only logarithmically, and *ϕ*_*c*_ decreases with Ω. The condition in Eq. (8) gives the lowest *ϕ* above which phage can drive bacteria below the extinction threshold *T* before bacteria resume growth (due to phage decay at rate *ω*). The leading factors determining the phage cocktail success are the initial rate of phage killing *ϕP*_0_ and bacteria growth *r*, which drive the transition to phage-bacteria coexistence. The dashed vertical green line in Figure 4A shows *ϕ*_*c*_ for Ω = 10. We observe that the numerical simulations agree with the heuristic prediction Eq. (8) when *I≪ I*_*b*_. Note that when phage are not very efficient (*ϕ≪ ϕ*_*c*_) the presence of resistance is not the major treatment failure driver, therefore Eq. (5) is an upper bound approximating the success condition. Together with the baseline single-phage treatment model results, our analysis of a simple phage cocktail model reveals that phage can restore therapeutic success even in immunodeficient hosts and even when phage-resistant bacterial mutants are present, insofar as sufficiently effective phage – according to the quantitative condition *ϕ > ϕ*_*c*_ – are utilized.

### C. Synergy between single-phage treatment and modulated responsive immune system against *in vivo* infections by multiple bacteria

The previous sections addressed the efficacy of monophage and phage cocktail therapy assuming that interactions between phage and bacteria occur in a well-mixed environment. Here we expand these results to a more realistic *in vivo* context by incorporating a nonlinear phage adsorption profile (reported to fit experimental adsorption curves [62, 63]), incorporating a structured phage infection dynamics, and including a dynamic response of the innate immune system. With the inclusion of these ingredients the model structure becomes similar to previous *in vivo* models that quantitatively reproduced experimental dynamics measured when treating lung infections with phage in immunomodulated mice [16]. Therefore we can directly compare our theoretical results with the parameters previously inferred, obtaining quantitative insights on the dynamics produced by single-phage treatments of *in vivo* infections.

Fig. 5 shows the final density of bacteria produced by numerical simulations of Eq. 3 after 200 hrs, for a broad range of immune system capacities *I* and phage adsorption rates *ϕ*. We find a qualitatively similar pattern as in Section III A, as the immune system needs to be strong enough to clear the infection in synergy with phage, which in turn can improve treatment outcomes with higher adsorption rates. We can compare infection outcomes for the parameter values explored in Fig. 5 to the parameters inferred in [16] (blue diamonds). Fig. 5 extrapolates the treatment outcome for biologically relevant parameters around these experimentally inferred values. Notably, we confirm that the current model predicts phage therapeutic success in immunocompetent mice (upper diamonds), however the model also predicts that this outcome is contingent on the use of a phage with a sufficiently rapid adsorption rate.

**FIG. 5:**
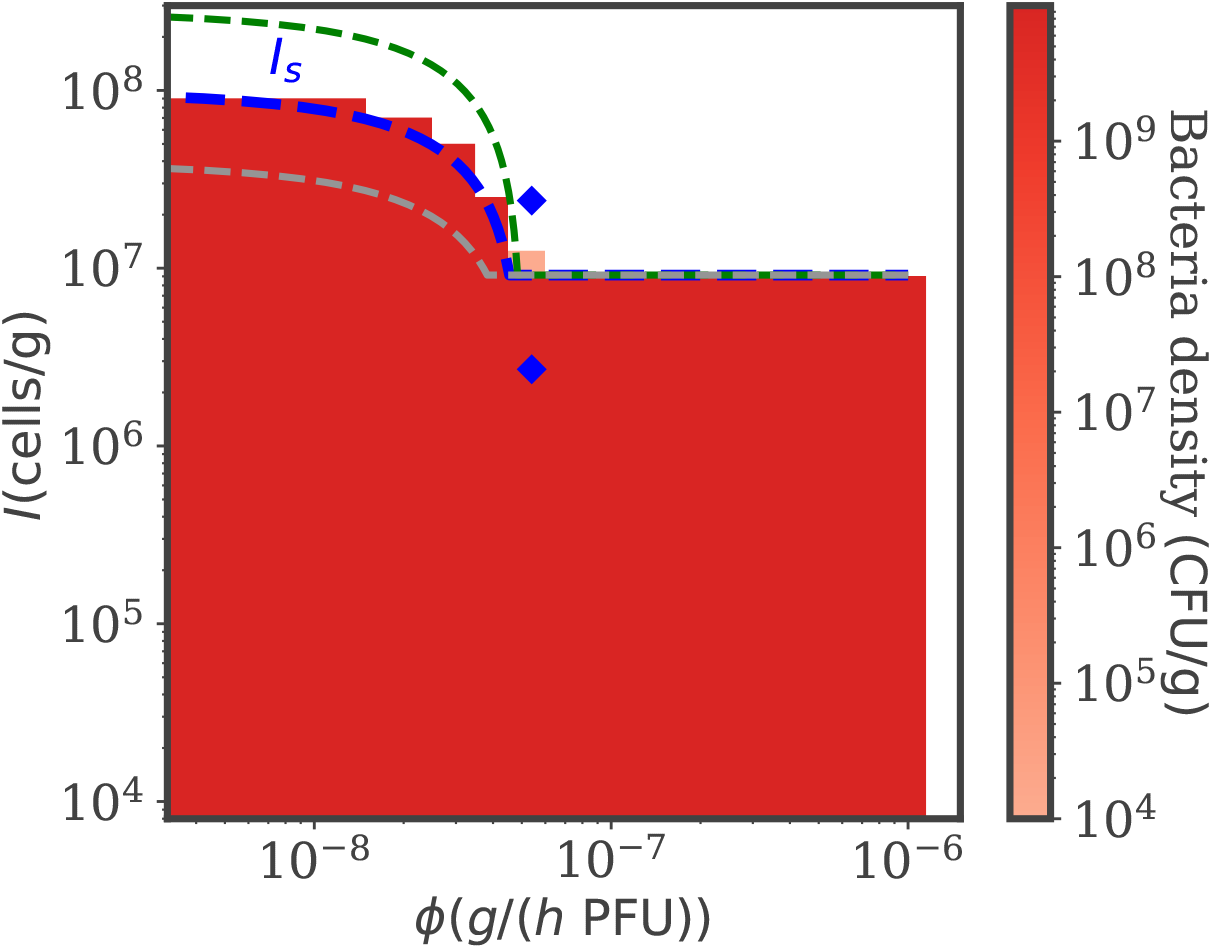
Single-phage therapy succeeds against phage resistance with a strong enough immune system. Numerical simulations of the model in Eq. (3) varying *I* and *ϕ*. The color map represents the density of bacteria in the last part of the numerical simulations. Single-phage therapy works with a strong enough immune system, which needs to be even stronger with inefficient phages (lower *ϕ*). The analytic condition in Eq. (9) (blue dashed line) predicts well the therapy outcome transition. The two blue diamonds correspond to the *I* and *ϕ* inferred in [16] from *in vivo* experiments on a model without infected bacteria classes. The green (grey) dashed line represents Eq. (9) with a 3-fold increase (decrease) in the concentration of bacteria at the beginning of the therapy. The bigger the bacteria inoculum (worse infection), the harder it is to clear the infection using phage with moderate to low *ϕ*. Simulation parameters are reported in Table II.

To understand these numerical results quantitatively, we proceed as in Section III A by considering the system without phage *P* = 0, which yields the same fixed points for bacteria density while the immune cells relax to *i*^*∗*^ = *I*. Hence Eq. (6) is again a necessary condition for therapy success, so that phage resistant bacteria can be controlled while still below the infectious dose 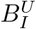. To progress further in the analysis we assume that phage infection dynamics are fast compared to other timescales, disregarding the dynamics of infected bacteria 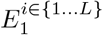, which corresponds exactly to the model in [16]. Assuming also a scenario in which phage are so abundant that *P≫ P*_*c*_, and that the immune cells dynamics relax fast to *I*, we find (details of the derivation in SI Sec. III D) a new immune threshold *I*_*S*_ that constrains the host immune capacity for which the single-phage therapy succeeds:

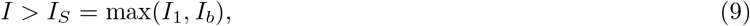

with

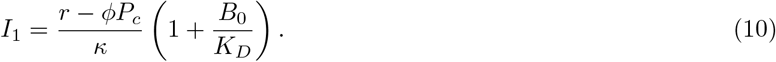

As in Section III A the condition for therapeutic success arises from combining two necessary immune constraints, so that phage and immune system can synergistically control both the susceptible bacteria (Eq. (10)), and the phage resistant mutants (Eq. (6)). Notably, the condition to overcome phage resistance in a timely fashion is determined by the balance between bacteria growth *r* and immune killing *κ* regardless of the new model ingredients. Eq. (9) depends on the infecting bacteria density at the time of therapy administration *B*_0_, highlighting the importance of prompt intervention when treating the infection. The dash blue line in Fig. 5 shows Eq. 9 as a function of *ϕ*. The analytical result agrees with the numerical results of the general model in Eq. 3 and with the findings of [16] where therapy worked in an immunocompetent cohort but failed in immunocompromised mice. The green and grey dashed lines in Fig. 5 show the quantitative impact of bacteria concentration at the treatment start, as they correspond to Eq. (9) with respectively a 3-fold increase and decrease in *B*_0_. We note that the highest *B*_0_ scenario brings the success to failure transition very close to the parameters that saw successful treatment in immunocompetent mice in [16].

### D. Robust benefits of therapeutic phage cocktails within an *in vivo* infection model, modulating immune responses and phage-bacteria interactions

Finally, we address the sensitivity of phage cocktail efficacy given variation in innate immune efficiency and phage life history traits. When adding a second phage to an *in vivo* infection model with a responsive modulated immune system and structured nonlinear phage-bacteria interactions, as described in Section II B 5, we predict that treatment would succeed if (see SI Sec. IV)

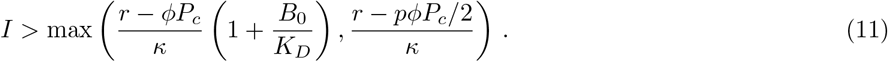

The first term in the maximum, the same as in Eq. (9), ensures that susceptible bacteria are killed, whereas the second term is the condition under which resistant types can be kept in check. Fig. 6 shows the infection clearance pattern as a function of *I* and *ϕ* for simulations of Model (4) with *p* = 1. The dashed lines show Eq. (11) for *p* = 0, 0.5 and 1 (blue, yellow, green). The black line represents a scenario where bacteria do not develop phage resistance, in which case the success condition is given just by Eq. (10). Our simulations agree with the analytical prediction in Eq. (11), as shown in *p* = 1 in Fig. 6 and in Fig. S1 for the other cases.

**FIG. 6:**
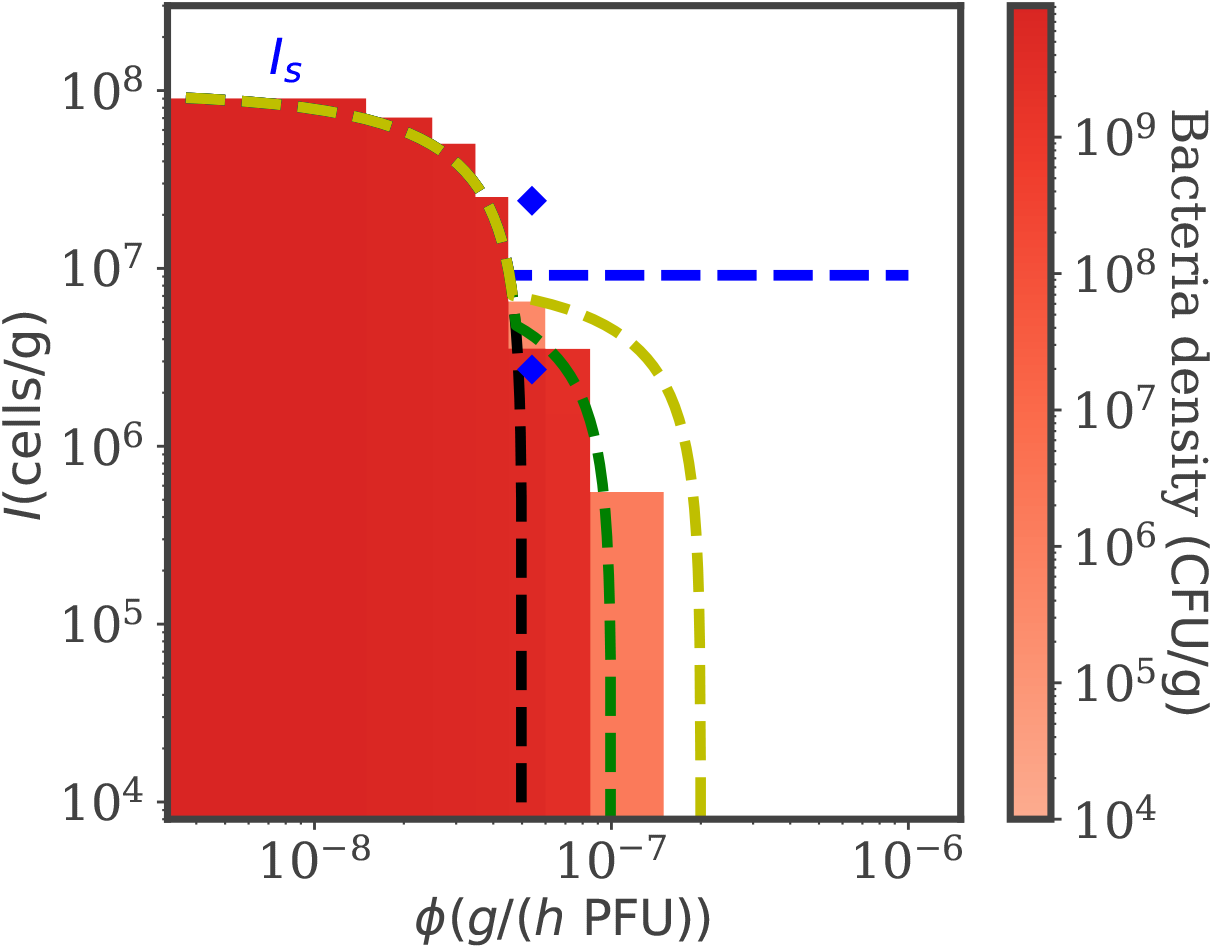
Efficient phage cocktails improve therapy success in an *in vivo* model of immunocompromised hosts. Numerical simulations of the model in Eq. (4) varying *I* and *ϕ*, with *p* = 1. The color map represents the density of bacteria in the last part of the numerical simulations. Phage cocktails can drive bacteria to extinction in immunocompromised hosts provided that phages have a high enough adsorption rate *ϕ*. The dashed lines show Eq. (11) for *p* = 0, 0.5 and 1 (blue, yellow, green), which agrees well with the simulation results. The black dashed line shows Eq. (10), representing therapy success when bacteria do not develop phage resistance. The two blue diamonds correspond to the *I* and *ϕ* inferred in [16] from *in vivo* experiments. Simulation parameters are reported in Table II.

The simulations findings in Fig. 6 show that the treatment with two phages improves the therapeutic outcome in immunocompromised hosts if the adsorption rate is high enough, confirming the qualitative picture presented in Section II B 3. Eq. (11) quantifies how the therapy success depends on phage-bacteria evolutionary interactions through variation in the cross-phage resistance parameter *p*. There is a critical 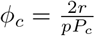 above which therapy succeeds when *I* = 0. We interpret this finding to mean that higher *p, i.e*. using therapeutic phages with more complementary phenotypes, improves the therapeutic outcome. In contrast, when *p* = 0 bacteria can evolve simultaneous resistance to both phages leading to the same result as in the single-phage treatment. Hence it is crucial to pre-select phages against which it is less likely that bacteria could evolve combined resistance, for instance making sure that they do not share a common receptor target, or that receptors targets are not pleiotropically linked [42], like in our LD experiment scoring a potential cocktail of PAK P1 and LUZ19v against *P. aeruginosa*. When *p* = 1 and the phage combination covers all bacteria resistant mutants, there is a region of parameters, between the black and the green lines in Fig. 6, where therapy fails due to phage resistance despite the phage cocktail treatment. Here the only way to improve the treatment efficacy would be to use phages with stronger lytic features.

Interestingly, when *p <* 2*/*3 and *P*_2_ is a generalist phage used alone, the maximum phage killing term in Eq. (4) against the most resistant bacteria type would be larger than when using two phages for the same value of *p* (see SI Sec. IV for the derivation). Therefore, it would be better to deliver *P*_2_ alone than together with another phage. Fig. S2 shows that, in the special case *p* = 0.5, a single *P*_2_ produces the same numerical results as a phage cocktail with *p* = 1, and indeed gives better outcomes than two phages with *p* = 0.5, as expected from the theory. This result is conditioned on the specific assumption we made in this model that phages compete to infect bacteria. Nevertheless it showcases an extreme situation where using a phage cocktail could be worse than a single phage, suggesting that it is crucial to quantify experimentally the effect of phage combinations in the system of interest. Even in this scenario, sub-optimal treatment with two phages with intermediate *p* still yields much better therapy outcomes in immunocompromised hosts than a treatment composed of a single or multiple phages that do not target all bacteria mutants in the pathogen population, as evident when comparing the dashed blue and yellow lines in Fig. 6. These results can be generalized to cases when the life history traits of the two phage differ from one another (see SI Sec. V). Altogether, this analysis reinforces the importance of selecting a combination of efficient phage against target bacteria when designing phage cocktails for treating infections in immunocompromised hosts.

## IV. DISCUSSION

In this work, analyzed simple tripartite population dynamic models of single-phage and phage combination treatments to clear infections by multiple strains of bacteria capable of evolving phage resistance in immunomodulated hosts. In doing so we extended previous work that considered the impacts of phage therapy when bacteria were exclusively susceptible to phage [26] by considering the potential combined use of phage with complementary modes of infection. This assumption is supported by new, fluctuation test experiments in which *P. aeruginosa* was unlikely to randomly acquire double resistance mutations to phage LUZ19v and PAK P1. Beginning with monophage treatment and extending this to phage cocktails, we find that immunophage synergy underlies the curative treatment of bacterial infections given sufficiently efficient phage. Notably, the use of phage cocktails can extend the range of immunocompromised conditions in which phage therapy can clear a pathogen. Finally, we extended core theoretical findings to a realistic *in vivo* modeling contexts, showing the robustness of immunophage synergy given variation in immune state, phage adsorption rates, and asymmetry in phage effectiveness within cocktails. Our analytical results quantify the importance of a prompt infection treatment and of selecting phages that are highly effective against the target pathogens including potential resistant mutants [65], especially when dealing with immunocompromised hosts. We extrapolate therapy outcome predictions around parameters inferred in *in vivo* lung infections by *P. aeruginosa* in immunomodulated mice [16], showing that moderate adsorption rate variations in any employed phage can have drastic effects on therapy outcomes, potentially making the difference between a successful and a failing treatment in experimental applications.

Our theoretical exploration of immunophage synergy builds on a set of assumptions that come with caveats to be addressed in future work - broadly speaking we categorize this in terms of simplifications in our representation of the immune system, phage infection and administration dynamics, and the evolutionary relationship between phage and bacteria. First of all, we consider a relatively simple impact of the host immune system on pathogens, through quantitative features that have been proposed in past infection models [16, 53, 59]. The first part of this study, focusing on non-responsive immunity is consistent with a signaling deficient immune system as in the case of myeloid differentiation primary response gene 88-deficient mice (MyD88^*−/−*^) [16]. The predictions in this limit may also be tested in *ex vivo* experiments mimicking infections in the lungs [66] by inoculating therapeutic phage and a fixed amount of immune cells. The innate immune responses considered in the second part of our work focuses on neutrophils, the first immune barrier against invading pathogens [67], while neglecting the other components of the innate immune response and the adaptive immune response altogether. Notably, inhibition of phage via immune responses [46] and by the reduction of circulating infectious phage by macrophages [25]. In the future it will be important to increase our quantitative understanding of the impact of the complex immune dynamics arising during infections.

The overall modeling structure used here adopts an implicit view of complex spatial processes taking place during phage treatment of respiratory lung infections [68]. Although it is possible to include effective, nonlinear interaction terms to mimic spatial complexity [16], moving forward it will be paramount to evaluate the explicit impact of spatial structure on quantitative phage-pathogens dynamics, for instance leveraging ex vivo technologies [69, 70], so that future models can incorporate spatial components [71]. In this study we also do not address the impact of treatment timing on therapy outcomes. A theoretical work applying control theory on a phage cocktail model suggested that additional treatment improvements may be possible by optimizing the timing and distribution of phage titers [72]. Such optimum would depend on the pathogen population composition and likelihood of resistance mutations, as well as on the quantitative features of phage-bacteria interactions during an infection, such as the functional shape of phage adsorption profiles. Simultaneous administration has been shown to outperform sequential treatments as controlling the pathogen population size right away reduces the chances of multi-resistance [40, 73], even though the generality of these results is unclear. In the future it will be important to integrate empirically constrained population dynamics models of local infections with pharmacokinetics parameters describing the likelihood and delay of delivering phage in the desired infected tissue [24].

Finally, analysis of phage cocktail impacts here assumes relatively simple evolutionary interactions between phage and bacteria such that bacteria cannot evolve resistance to both phages at the same time. Our choice is motivated by previous works that suggested combinations of phages that target different bacteria receptors in order to improve treatment efficacy [41, 42, 54]. This hypothesis is supported by fluctuation test findings in which *P. aeruginosa* did not randomly develop simultaneous resistance to LUZ19v and PAK P1 (see SI Section III). It is important to note that although a wild type population susceptible to both phages need not necessarily evolve a double resistant mutant immediately, this does not preclude the potential for a population already selected to persist given exposure to one of the two phages may evolve to become double resistant. The current study shows that phage cocktails can perform robustly even when modulating rules regarding phage complementarity. In the future it will be crucial to further explore more complex eco-evolutionary processes that can arise between pathogenic bacteria and therapeutic phage during the course of an infection, whether in an acute or chronic infection context.

Despite these caveats, the combined use of experiments, simulations and theory provide guidance on the expected range of phage therapeutic efficacy whether using monophage or phage cocktail treatments. Building upon earlier findings [16], our framework provides testable predictions on the quantitative impact of different modes of tripartite phage-bacteria-immune interactions on therapeutic outcomes over a range of host immune conditions and phage life history traits, highlighting the success of single-phage therapy in synergy with a strong enough immune system and the benefit of phage combination therapy in immunocompromised hosts. Importantly, the parameters inferred in [16] fall right at the boundary between treatment failure and success in immunodeficient hosts, which could result in drastic differences in the infection outcomes given small variations in phage and immune features. Such sensitivity makes it crucial to inform model development with *in vitro* and *in vivo* data to improve therapeutic design.

More broadly, this work represents a further step towards quantitatively addressing the impact of evolutionary considerations on phage therapy outcomes. Here we showcased a specific example of how we can harness the evolutionary potential of phages to develop phage cocktails that target specific strains of multi-drug resistant bacteria, facing the challenge posed by the evolution of phage resistance [42, 43]. Whether we seek to exploit *in vitro* phage training [30, 43] or evolutionary trade-offs [31–36], in the future it will be essential to integrate further experimental evidence into quantitative models, tackling different aspects of phage-bacteria evolutionary interactions during therapy. Predictive models that integrate the tripartite interactions among pathogenic bacteria, therapeutic phages, and the eukaryotic host, along with evolutionary processes, will be crucial for designing effective phage treatment strategies both in the near and long term.

## Acknowledgements

The work was supported by the National Institutes of Health (R01 AI146592 to JSW, LD). JSW was supported, in part, by the Chaires Blaise Pascal program of the Ile-de-France region. We thank Jeremy Seurat and Rogelio Rodriguez-Gonzalez for insights in the development of the *in vivo* phage therapy model.

