## Supplementary Information for "Multi-strain phage induced clearance of bacterial infections"

### Supplemental Information: Multi-strain phage induced clearance of bacterial infections

#### I. BACTERIA AND PHAGE STRAINS FOR THE FLUCTUATION TEST

The *P. aeruginosa* strain PAK-lumi[74] was grown in semi-solid or in liquid (with shaking) lysogeny broth Lennox medium (Difco) at 37°C. The virulent phages PAK_P1 (93kb, NC_015294.2)[75] and LUZ19v (43kb) [76] were amplified on strain PAK-lumi. Large lysates (500 mL) were concentrated by ultrafiltration and subsequently ultracentrifuged twice using CsCl gradient [77]. The concentrated phage suspension was dialyzed against Tris 10 mM pH7 NaCl 200 mM and store at 4°C until use.

#### II. FLUCTUATION TEST

A single colony of strain PAK-lumi was picked from a Lennox plate to inoculate 5mL of LB Lennox liquid broth and incubated at 37°C with shaking (180 rpm) during 1 hour. Then the OD at 600nm was adjusted with fresh medium to reach 0.1 and subsequently diluted in LB broth to obtain $5 \cdot 10^3$ CFU/mL in 10mL. A sample of 50µL was withdrawn and spreaded on a LB agar plate and incubate at 37°C overnight before counting CFUs. Next, the diluted culture was used to fill 60 wells of a sterile 96 well flat-bottom microplate (Falcon, Cat Id 353072) with 100µL per well. The microplate was covered with a clear film and wrapped in parafilm to prevent evaporation before overnight incubation with shaking at 37°C. The next day, the bioluminescence emitted from each well was recorded using GloMax Discover Microplate Reader (Promega). Then, in 10 wells we added 100µL of phosphate buffered saline (PBS) (Sigma-Aldrich) while in the fifty other wells we added 100µL of $10^9$ plaque-forming unit/mL (PFU/mL) of phage PAK_P1 or LUZ19v or their mixture (50% of each). The microplate was incubated at 37°C with shaking at 180rpm for 3 hours. Next, we diluted each of fifty wells by 10 (20µL of each well introduced into another plate containing 180µL of PBS) and spread100µL on LB agar plates that were previously overlayed with 1mL of $10^7$ PFU/mL of phage PAK_P1 or LUZ19v or their mixture (50% of each) [3]. The 10 remaining wells (no phage) were used to determine the number of colonies by plating serial dilutions on LB agar plates. All agar plates were incubated overnight at 37°C before counting CFUs. We performed these tests six times independently for each of the two phages and three times for the mixture.

#### III. MUTATION PROBABILITIES INFERENCE

From the count of PAK-lumi mutants resistant to either phage PAK_P1, LUZ19v or both we infer the probability at which cells develop resistance to the corresponding phage (or their combination) within one duplication. Under the null assumption that mutations are random [3], the number of mutants follows the LD distribution [78]. We run the inference through the web-tool bz-rates [44], which learns the model parameters via the Generating Function estimator from [45], and allows for differential growth rates and a plating efficiency lower than 1. According to the protocol described in the previous section in which the plated volume is 1/20 the original well volume, we use a plating efficiency of 0.05, after dividing the mutants and control counts by 8 to correct for the excess growth during the 3 hours incubation time after the overnight growth. The LD distribution passes the $\chi^2$ goodness-of-fit test against the experimentally measured distributions for the number of mutants, for both phages. Through this procedure, we find that *P. aeruginosa* develop resistance to PAK_P1 with a probability $3.7 \cdot 10^{-8}$ per duplication (CL between $3.4 \cdot 10^{-8}$ and $4 \cdot 10^{-8}$). Resistance to LUZ19v emerges at a probability $4.9 \cdot 10^{-8}$ per duplication (CL between $4.5 \cdot 10^{-8}$ and $5.3 \cdot 10^{-8}$). For sake of simplicity, in the models presented in this work we assume that bacteria develop resistance to both phages with the same probability $4.3 \cdot 10^{-8}$ per duplication, which is the average of each phage mutation probability and is reasonably close to the Confidence Levels bounds for both phages. We don't observe a single double-resistant mutant in any of the 150 colonies plated against the phage cocktail.

##### A. Numerical implementation

We simulate the models introduced in Section II via a Python ODE integrator available in the Scipy library [79], with an explicit Runge-Kutta method of order 5(4) [80]. The model parameters are set to the values indicated in Tables I and II respectively.

For the models (1) and (2) we start with $P_0 = 4K_C = 4\frac{r}{d}$ phage, and then to avoid singular initial conditions we constrain $P_0$ between 100 PFU/g and $10P^*$, with the resistance-free equilibrium phage density being $P^* = \frac{r}{\phi}\left(1 - \frac{\omega}{\beta\phi K_C}\right) - \frac{\kappa I}{\phi + \frac{\omega}{\beta K_D}}$. Then, when implementing a cocktail treatment, each phage is initialized at a density $\frac{P_0}{2}$. Susceptible bacteria start at a density $S_0 = \frac{K_C}{10} = \frac{r}{10\tilde{d}}$, while also being constrained between 100 CFU/g and $10S^*$, where the resistance-free equilibrium bacteria density reads $S^* = \frac{\omega}{\beta\phi}$. Finally the initial density of bacteria resistant to any of the present phage is set to $R_0 = 10$ CFU/g. For the models (3) and (4) we initialize the simulations with values close both to those inferred in [16] and presented in [26]. Therefore we start with $P_0 = 10^7$ PFU/g, $S_0 = 4 \cdot 10^8$ CFU/g and $R_0 = 10$ CFU/g. All intermediate infected bacteria states are initialized to 0 CFU/g. In this case the initial immune cells density is set to the minimum between $2.7 \cdot 10^6$ neutrophils/g and $\frac{I}{5}$.

We run the simulations up to 200 hours, while stopping the simulations when the infection is cleared, that is when the total bacteria density $B_{tot}$ goes below 10 CFU/g corresponding to about 5 bacteria in 0.5 g of infected tissue (about the weight of mice lungs). Once the simulation is completed we score the efficacy of phage treatment by computing the average density of bacteria in the last 24 hours, unless bacteria go extinct before then in which case the number of bacteria is set to 0 (treatment success).

The source code is available at `https://github.com/Jacopo-Marchi/multi-strain-infection-clearance`

##### B. Positive infectious dose threshold

If there are no phages ($P = 0$) we consider the dynamics of the total amount of bacteria $B = S + R$ for the model introduced in II B 2:

$$\frac{dB(t)}{dt} = rB - \tilde{d}B^2 - \frac{\kappa IB}{1 + \frac{B}{K_D}}. \tag{S1}$$

As shown in [26] the stability analysis yields an unstable fix point at

$$B_I^U = \frac{1}{2}(\frac{r}{\tilde{d}} - K_D) - \Delta \tag{S2}$$

and a stable one at

$$B_I^S = \frac{1}{2}(\frac{r}{\tilde{d}} - K_D) + \Delta, \tag{S3}$$

with $\Delta = \sqrt{(\frac{r}{\tilde{d}} + K_D)^2/4 - \frac{r}{\tilde{d}}K_D\kappa I/r}$. The model also permits a fixed point at $B^* = 0$ . Importantly, this fixed point is stable if $B_I^U > 0$, which is the necessary condition for therapy success in presence of phage resistance, presented in the main text.

There is an immune system threshold $I > I^c$ above which the immune system always clears the infection on its own and the only stable fix point is $B = 0$. From Eq. (S1) the condition for $I^c$ is given by $\frac{\kappa I^c B}{1 + \frac{B}{K_D}} > rB - \tilde{d}B^2$, $\forall B$. We have:

$$I^c = \frac{r}{4\kappa}\left(\sqrt{\frac{r}{\tilde{d}K_D}} + \sqrt{\frac{\tilde{d}K_D}{r}}\right)^2. \tag{S4}$$

If $K_D \ll r/\tilde{d}$, we have $\Delta \sim \frac{r}{2\tilde{d}}\sqrt{1 - \frac{\kappa I}{r}\frac{4K_D\tilde{d}}{r}}$ and

$$I^c \sim \frac{r}{4\kappa}\frac{r}{K_D\tilde{d}}. \tag{S5}$$

For $I \ll I^c$ we have $\Delta \sim \frac{r}{2\tilde{d}}(1 - \frac{\kappa I}{r}\frac{2K_D\tilde{d}}{r})$ and

$$B_I^U \sim K_D(\frac{\kappa I}{r} - 1). \tag{S6}$$

Under these assumptions the condition $B_I^U > 0$ reads Eq. (6). Note that the two conditions $K_D \ll r/\tilde{d}$ and $I_b \ll I^c$ are consistent with one another, as evident from combining Eqs. (S5) and (6).

##### C. Heuristic derivation for the cocktail therapy success condition

When using a therapy composed of two phages, bacteria can be susceptible to either one of the phages or to both, according to the model (2). The infection can be cleared even in completely immunocompromised hosts ($I = 0$), if $\phi > \phi_c$ where $\phi_c$ satisfies the transcendental equation (8).

To derive this condition, consider a scenario in which therapy is successful and bacteria are driven to extinction. Eventually $S \ll \frac{\omega}{\beta\phi}$ and phages will decay exponentially at rate $\omega$, as $P_1(t) = P_2(t) = P(t) = P_0 e^{-\omega t}$. In this case we also have $B(t) \ll \frac{r}{\tilde{d}}$. The dynamics for the resistant types ($R_1 = R_2 = R$, since the system is symmetric) in the $I = 0$ limit then read

$$\frac{dR(t)}{dt} = rR - R\phi P_0 e^{-\omega t} \,. \tag{S7}$$

Therefore resistant types will decrease if $1 > e^{-\omega t} > \frac{r}{\phi P_0}$, or

$$t < t_R = -\frac{1}{\omega} \ln\left(\frac{r}{\phi P_0}\right) . \tag{S8}$$

Similarly, the dynamics for the susceptible bacteria are given by

$$\frac{dS(t)}{dt} = rS - S2\phi P_0 e^{-\omega t} \,. \tag{S9}$$

The time for which $\dot{S} > 0$ is $t_S = t_R + (\ln 2)/\omega$, which is bigger than $t_R$ . We can integrate by parts to find the time $t_E$ at which bacteria decrease below the extinction threshold $T$, starting from $S = \frac{\omega}{\beta\phi\Omega}$, where $\Omega \gg 1$ is a numerical factor ensuring our initial assumption of $S \ll \frac{\omega}{\beta\phi}$, while $P \sim P_0$. Solving for $S(t_E) = T$ gives $\ln \frac{T\beta\phi\Omega}{\omega} = rt_E - \frac{2\phi P_0}{\omega}(1 - e^{-\omega t_E})$, which can be rewritten as:

$$\frac{1}{r} \ln \frac{T\beta\phi\Omega}{\omega} + \frac{2\phi P_0}{\omega r} = t_E + \frac{2\phi P_0}{\omega r} e^{-\omega t_E} := F(t_E) \,. \tag{S10}$$

Therapy succeeds if bacteria go extinct before $R$ start regrowing, $t_E < t_R$. We define the function $F(\cdot)$ as the right-hand side of Eq. (S10). $F' < 0$ for $t_E < t_S$, hence the therapy success condition is $\frac{1}{r} \ln \frac{T\beta\phi\Omega}{\omega} + \frac{2\phi P_0}{\omega r} = F(t_E) > F(t_R)$. Plugging in Eq. (S8) we get

$$\frac{2\phi P_0}{r} > 2 + \frac{\omega}{r} \ln \frac{\omega}{T\beta\phi\Omega} + \ln \frac{\phi P_0}{r} \,, \tag{S11}$$

which leads to (8). Note that, for this argument to be valid, we need the extra constraint that phage decay (below $P_0$) starts before bacteria extinction: $\frac{\omega}{\beta\phi\Omega} > T$. If there is no such $\Omega$, the condition $\frac{r}{\phi P_0} < 1$ will be sufficient for bacteria extinction.

We solve the transcendental equation recursively plugging in $P_0$ (see Sec. III A) and updating the value of $\phi$ at each iteration according to (S11), starting from the lower bound $\frac{r}{P_0}$. We confirmed the result with the Newton-Raphson algorithm [81] which converges to the same value.

##### D. Single-phage therapy success condition in a realistic infection model

To derive the therapy success condition from (3), we assume that phage infection dynamics are fast, removing infected bacteria $E_1^{i\in\{1...L\}}$, as in [16]. We also assume that phage are so abundant that $P \gg P_c$, and that the immune cells dynamics relax fast to $I$. Under these assumptions, in the absence of phage resistance the only slow variable are the susceptible bacteria $S$, whose equation reads

$$\frac{dS(t)}{dt} = rS - \tilde{d}S^2 - \frac{\kappa IS}{1 + \frac{S}{K_D}} - S\phi P_c. \tag{S12}$$

If $r - \phi P_c < 0$, $S$ goes to 0, but therapy still requires Eq. (6) to succeed in light of phage resistance. More precisely, this equation does not admit real nonzero fix points if $I > \tilde{I}_c = (\frac{r-\phi P_c}{\tilde{d}} + K_D)^2 \frac{\tilde{d}}{4K_D\kappa}$. If $\frac{r-\phi P_c}{\tilde{d}} \leq CK_D$ with $C \sim O(1)$

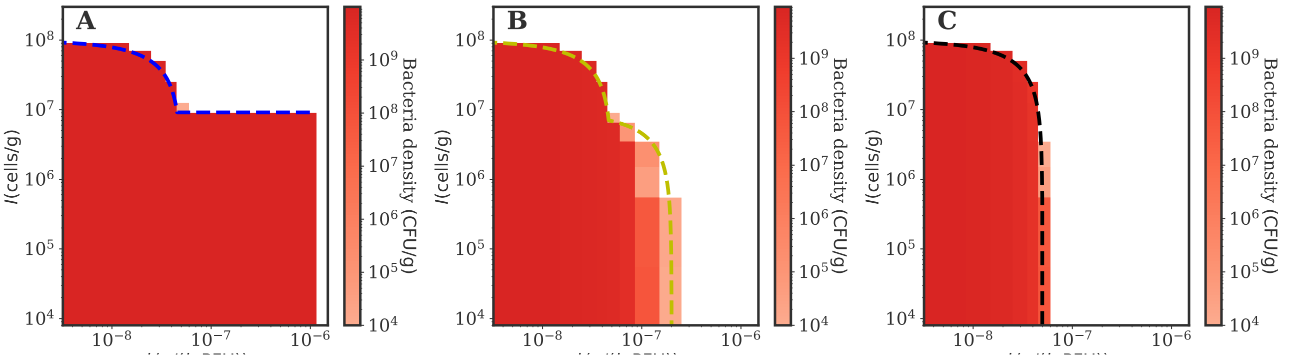


FIG. S1: **Numerical simulations confirm the cocktail success condition** Numerical simulations of the model in Eq. (4) varying $I$ and $\phi$, with $p = 0$ (A), $p = 0.5$ (B) and without allowing for phage resistance (C). The colormap represents the density of bacteria in the last part of the numerical simulations. The dashed line represents the prediction in Eq. (11) for $p = 0$ in A) and 0.5 in B) (blue and yellow respectively). The black dashed line in C) shows Eq. (10), representing therapy success when bacteria do not develop phage resistance. Simulation parameters are reported in Table II.

we have $I_c \leq (C+1)^2 \frac{K_D \tilde{d}}{4\kappa} \ll I_b$, where we used $K_D \ll r/\tilde{d}$. Hence Eq. (6) is a sufficient condition for infection clearance.

Conversely, in the limit $\frac{r-\phi P_c}{\tilde{d}} \gg K_D$, for $I < \tilde{I}_c \sim \frac{r-\phi P_c}{\tilde{d}} \frac{r-\phi P_c}{4K_D\kappa}$, we have a pair of stable and unstable fix points for $S$, $S^S_{I,P}$ and $S^U_{I,P}$ respectively, that can be obtained simply by substituting $r$ with $r - \phi P_c$ in Eqs. (S2) and (S3). If $I \ll \tilde{I}_c$, we get that $S^U_{I,P} > B_0$ if

$$I > I_1 = \frac{r - \phi P_c}{\kappa}\left(1 + \frac{B_0}{K_D}\right) , \tag{S13}$$

which combined with Eq. (6) gives the condition for single phage therapy success Eq. (9) in the main text.

As a side note, the self-consistency condition $I_1 \ll \tilde{I}_c$ breaks if $\frac{r-\phi P_c}{\tilde{d}} \sim O(B_0)$, but for the parameters extracted from [16] for such values of $\phi P_c$, $\tilde{I}_c \sim O(I_b)$, in which case therapy would succeed due to Eq. (6). Therefore our approximation is relevant to study the therapy success transition in this work.

Furthermore, we can prove that $B^U_I < S^S_{I,P}$ and therefore this condition does not affect therapy success. In the relevant limit $\frac{r-\phi P_c}{\tilde{d}} \gg K_D$ the condition reads $\frac{K_D\kappa I}{r} - \frac{r-\phi P_c}{2\tilde{d}} > \sqrt{\frac{r-\phi P_c}{2\tilde{d}}^2 - \frac{K_D\kappa I}{\tilde{d}}}$. The left-hand side is $> 0$ only if $I > 2\tilde{I}_c \frac{r}{r-\phi P_c}$, in which case $S$ has no other fix point than 0.

#### IV. PHAGE COCKTAIL SUCCESS IN A REALISTIC INFECTION MODEL

In order to derive the success condition for a phage cocktail described by model (4), we assume, as in the previous section, that phage infection dynamics are fast, and that phage eventually satisfy $P_1, P_2 \gg P_c$, therefore $F(P_1) + F(P_2) \to P_c$. Again, these assumptions lead to the condition (S13) that susceptible bacteria can be wiped out.

In addition to Eq. (S13) we need to derive the condition under which resistant bacteria can be kept in check. This translates to the condition $\dot{R}_1 < 0$, since in our model $R_1$ is harder to control as it is only killed by $P_2$. We assume that in the beginning of the simulations the dynamics of $P_1$ and $P_2$ are equivalent since phage replication is mainly driven by susceptible bacteria that are infected equally by both phages. Therefore soon after treatment we have $P_1 \sim P_2 \gg P_c$, and $F(P) \to P_c/2$. Note that the approximation $P_1 \sim P_2$ holds exactly if $p = 1$ since in this case the equations are symmetric with respect to the phage index. If therapy is successful, we can assume that at some point $B_{tot} \ll K_D$. Combining these assumptions we have that $\dot{R}_1 \sim rR_1 - \kappa I R_1 - pR_1\phi P_c/2$. Therefore resistant bacteria die out if

$$I > I_2 = \frac{r - p\phi P_c/2}{\kappa} , \tag{S14}$$

which combined with Eq. (S13) yields the cocktail therapy success condition in Eq. (11). Fig. S1 verifies the theoretical prediction against numerical simulations, showing that it holds for different values of $p$. In Panel C we do not include

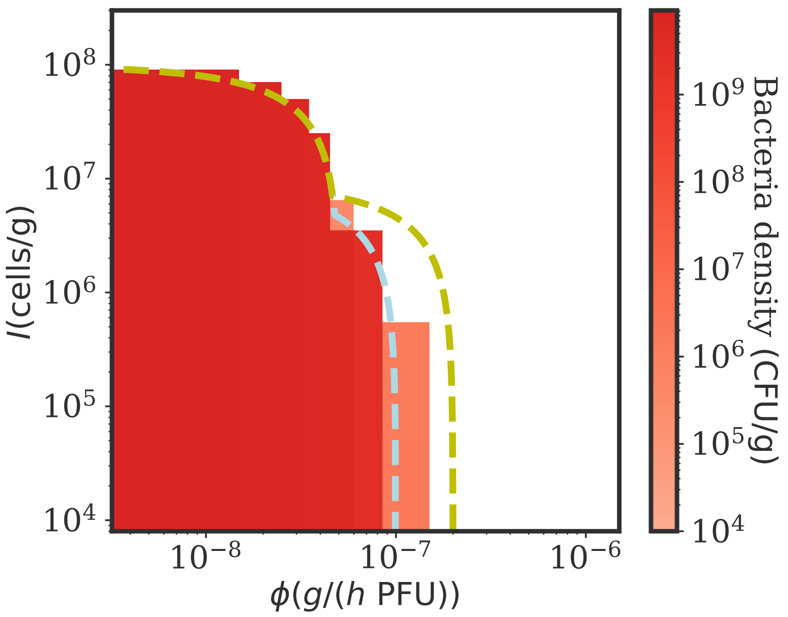


FIG. S2: **A generalist phage works better alone than in combination** Numerical simulations of the model in Eq. (4) varying $I$ and $\phi$, with a single phage $P_2$ and $p = 0.5$. The colormap represents the density of bacteria in the last part of the numerical simulations. In this extreme case, a single-phage treatment works better than two phages. The dashed lines compare Eq. (11) for $p = 0.5$ with two phages (yellow) with the transition for a single generalist phage (lightblue, equivalent to Eq. (11) with $p = 1$ ). Simulation parameters are reported in Table II.

resistant types, in which case the only condition necessary for successful treatment is the elimination of $S$, denoted by Eq. (S13).

As highlighted in Section III D, the maximum phage killing term for $R_1$ in Eq. (4) with two phages, always smaller than the phage killing of $R_2$, satisfies $\tilde{D} = p\phi F(P_2) \to p\phi P_c \frac{P_2}{P_1+P_2} \sim p\phi P_c/2$. With just the phage $P_2$, this term reads $D_1 = p\phi F(P_2) \to p\phi P_c$, whereas for $R_2$ we have $D_2 = (1-p)\phi F(P_2) \to (1-p)\phi P_c$ . Given our modeling choices of the cross-infection network and competition among phage types through $F(P)$, there are some intermediate values of $p$ for which $\tilde{D} < \min(D_1, D_2)$. This is always true for $p < 0.5$ since $\tilde{D} < D_1$. $\tilde{D} < D_2$ implies $p < 2/3$. Therefore, it is better to use a generalist phage $P_2$ (intermediate values of $p$) on its own rather than together with other phages. In particular, if $p = 0.5$ and $P_2$ is a generalist used alone, the maximum phage killing term in Eq. (4) would be the same as in the case of two phages with $p = 1$. Fig. S2 shows that indeed a single $p = 0.5$ phage produces the same numerical results as a combination when $p = 1$, and gives better outcomes than two phages with $p = 0.5$.

#### V. PHAGE COCKTAILS WITH DIFFERENT PHAGE LIFE TRAITS

Finally, we relaxed the assumption that different phages have the same life traits (assuming $p = 1$). We set $\phi_2 = \sigma\phi_1$ so that the adsorption rate of the second phage $P_2$ is different than that of $P_1$. As a result, therapeutic phage and pathogenic bacteria strains engage in coevolutionary dynamics during the infection with turnovers in the population dominant type (see Fig. S4 for an example), eventually producing extinctions (therapy success) or coexistence (therapy failure). Fig. S3 shows the resulting infection clearance pattern as a function of $I$ and $\phi_1$ for $\sigma = 1/5$ (A) and $\sigma = 5$ (B). As we can see, if the second phage has a different adsorption rate the therapy outcome can differ drastically, with worse phages severely worsening treatment in immunocompromised cases (A), and better phages improving the outcome overall with a stronger impact in immunocompetent subjects (B). Comparing the outcomes in Fig. S3 with the parameters extracted from [16] (blue diamonds), we note that the adsorption rate of any of the therapeutic phages may have drastic effects on the experimental outcome of phage therapy. An inefficient second phage (panel A) jeopardizes treatment success in immunocompromized hosts even if the first phage was much more lytic than the one used in [16]. On the other hand, an efficient phage addition (panel B) enhances the benefits of phage cocktails almost pushing the success condition beyond the immunocompromised experiment that failed in [16].

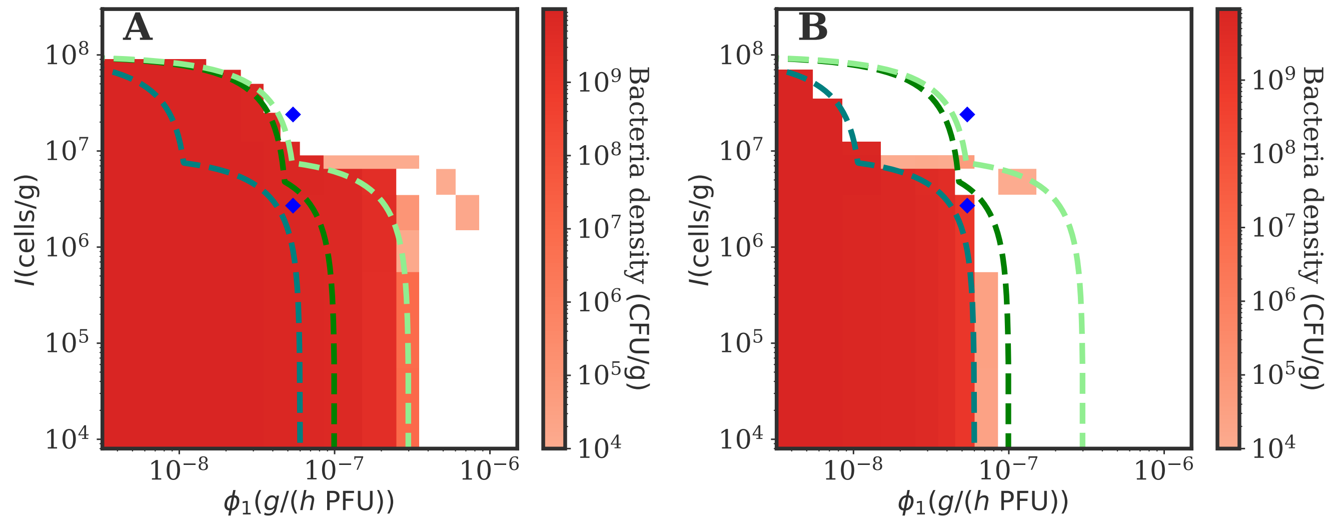


FIG. S3: **Phage cocktail treatments with different viral traits produce different outcomes.** Numerical simulations of the model in Eq. (4) with $\phi_2 = \sigma\phi_1$, varying $I$ and $\phi_1$, for $\sigma = 1/5$ (A) and $\sigma = 5$ (B). The color map represents the density of bacteria in the last part of the numerical simulations. A) While phage cocktails can still drive bacteria to extinction in immunocompromised hosts, a low $\sigma$ can jeopardize the treatment outcome. B) A high $\sigma$ improves therapy, especially in immunocompetent hosts. The dashed lines show Eq. (S19) for $\sigma = 1/5$, 1 and 5 (light green, green, teal). The simulations agree reasonably well with the analytical approximation. The two blue diamonds correspond to the $I$ and $\phi$ inferred in [16] from *in vivo* experiments. Simulation parameters are reported in Table II.

If $\phi_2 = \sigma\phi_1$ the system is not symmetric anymore with respect to phage index, since from the beginning the different phage types replicate at a different rate through the susceptible bacteria. Therefore, when $P_1, P_2 \gg P_c$, we cannot assume that $F(P) \to P_c/2$. Instead we have $F(P_1) \to P_c\frac{P_1}{P_1+P_2}$ and $F(P_2) \to P_c\frac{P_2}{P_1+P_2}$ with a phage effect that is modulated by the relative frequency of the corresponding phage type. If the phage abundances were still symmetric, the success condition would simply read

$$I \sim \max\left(\frac{r - \phi_1 P_c \frac{1+\sigma}{2}}{\kappa}\left(1 + \frac{B_0}{K_D}\right), \frac{r - \phi_1 P_c \frac{\min(1,\sigma)}{2}}{\kappa}\right), \tag{S15}$$

where the first term corresponds to the first term in Eq. (11) taking the average $\phi$, and the second term is the same upon taking the minimum $\phi$ so that the target of the less lytic phage is cleared. Here we provide a phenomenological, back of the envelope argument to suggest a first order correction that accounts for the imbalance in phage densities. The equation for the time evolution of $\frac{P_1}{P_1+P_2}$ does not allow detailed analytical treatment, but with this section we provide some intuition on how the success condition depends on phage asymmetry $\sigma$.

Figure S4 shows an example of some dynamics for $\sigma = 1/5$, $\phi_1 = 5 \cdot 10^{-7}$ and low immune capacity ($I = 10^4$), where the simulations indicate that the infection is cleared while Eq. (S15) would predict therapy failure. In a first part of the dynamics phages grow on the susceptible types based on the corresponding adsorption rate, till $P_1, P_2 \gg P_c$ (or until the infection is cleared). After the first fast phage growth (first peak in $P_1$ and $P_2$ in Figure S4), the condition that $S$ is killed reads

$$I > \frac{r - P_c(\phi_1 \frac{P_1}{P_1+P_2} + \phi_2 \frac{P_2}{P_1+P_2})}{\kappa}\left(1 + \frac{B_0}{K_D}\right), \tag{S16}$$

in which the different adsorption rates contribute through the weighted average via the phage frequencies. Since the phage with the highest $\phi$ is abundant, this success condition is always lower than what predicted by Eq. (10). This approximation can be corrected, to first order, accounting for the excess of phages produced within a lysis event , for which $P_1/P_2 = \beta\phi_1 S_0/\beta\phi_2 S_0$, hence $P_1/(P_1 + P_2) = \phi_1/(\phi_1 + \phi_2)$. This yields

$$I > \frac{r - \phi_1 P_c \frac{1+\sigma^2}{1+\sigma}}{\kappa}\left(1 + \frac{B_0}{K_D}\right). \tag{S17}$$

Indeed in Figure S4 we see that $P_1$ reaches a higher peak than $P_2$, which causes bacteria susceptible to the more lytic phage (in this case $S$ and $R_2$, as we assume $\phi_1 > \phi_2$ without loss of generality) to be cleared, until bacteria

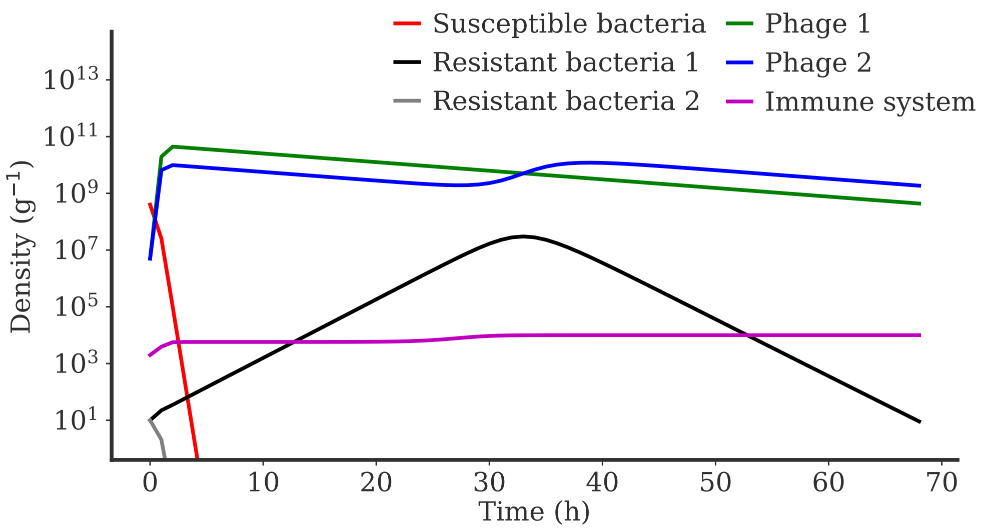


FIG. S4: **Simulation dynamics with different $\phi$s.** Numerical simulations of the model in Eq. (4) varying $I$ and $\phi_1$, for $\sigma = 1/5$ $\phi_1 = 5 \cdot 10^{-7}$ and $I = 10^4$. The rest of the simulation parameters are reported in Table II.

population is dominated by the target of the less lytic phage (the bacteria population turns over to $R_1$). In a second dynamical phase (also visible in the example dynamics in Figure S4 ), $P_1$ decays at a rate $\omega$. $R_1$ grows till triggering a burst of $P_2$ above $P_1$. We assume that $R_1$ can finally be cleared when the relative phage abundances swap so that $P_2/(P_1 + P_2) \sim \phi_1/(\phi_1 + \phi_2)$. When this population turnover is realized, the death terms of bacteria $R_1$ exceed growth, leading to clearance, if

$$I > \frac{r - \phi_2 P_c \frac{\phi_1}{\phi_1+\phi_2}}{\kappa} = \frac{r - \phi_1 P_c \frac{\sigma}{1+\sigma}}{\kappa} . \tag{S18}$$

Combining Eqs. (S17) and (S18) we obtain the condition under which both resistant types can be controlled

$$I \sim \max\left( \frac{r - \phi_1 P_c \frac{1+\sigma^2}{1+\sigma}}{\kappa} \left(1 + \frac{B_0}{K_D}\right), \frac{r - \phi_1 P_c \frac{\sigma}{1+\sigma}}{\kappa} \right) . \tag{S19}$$

Therefore, when $I = 0$, the infection is cleared if $\phi_1$ is bigger than $\phi_c = \frac{r(1+\sigma)}{\sigma P_c}$. A treatment with a worse second phage ($\sigma < 1$) is better than with only one phage since it can still clear the infection if the first phage is really efficient. On the other hand we see that a lower $\phi_2$ can dramatically compromise the treatment in immunocompromised hosts. When one of the two phages gets really good (high $\sigma$), the success transition saturates at $\phi_c \to \frac{r}{P_c}$, which is the limit of success when resistance does not emerge in the $\sigma = 1$ case. The dashed lines in Fig. S3 show Eq. (S19) for $\sigma = 1/5$, 1 and 5 (light green, green, teal). The simulations agree reasonably well with the back of the envelope estimate.
